# Drivers of nematode diversity in forest soils across climatic zones

**DOI:** 10.1101/2022.03.18.484848

**Authors:** Yuanhu Shao, Zuyan Wang, Tao Liu, Paul Kardol, Chengen Ma, Yonghong Hu, Yang Cui, Cancan Zhao, Weixin Zhang, Dali Guo, Shenglei Fu

## Abstract

Nematodes are the most abundant multi-cellular animals in soil, influencing key processes and functions in terrestrial ecosystems. Yet, little is known about the drivers of nematode abundance and diversity in forest soils across climatic zones. This is despite forests cover approximately 30% of the earth’s land surface, provide many crucial ecosystem services but strongly vary in hydrothermal conditions and associated ecosystem properties across climatic zones.

Here, we collected nematode samples from 13 forests across a latitudinal gradient. We divided this gradient in temperate, warm-temperate, and tropical climatic zones. Using boosted regression trees, we showed that across the gradient, nematode abundance and diversity were mainly influenced by soil organic carbon. However, within climatic zones, other factors were more important in driving nematode alpha-diversity, nematode biomass and the abundance of different trophic groups: mean annual temperature and total soil phosphorus in temperate zones, soil pH in warm-temperate zones, and mean annual precipitation in tropical zones. Additionally, nematode beta-diversity was higher in temperate than in warm-temperate and tropical zones, and we did not find significant differences among climatic zones in nematode gamma diversity.

Together, our findings indicate a latitudinal shift in the main climatic variables controlling soil nematode communities and demonstrate that the drivers of soil nematode diversity in forested ecosystems are affected by the spatial scale and climatic conditions considered. This implies that high resolution studies are needed to accurately predict how soil functions respond if climate conditions move beyond the coping range of soil organisms. Our results also emphasize the importance of studying the area-diversity relationship of soil organisms under different climatic conditions.

## Introduction

Soil biodiversity accounts for a large fraction of the biodiversity on Earth and is key for the functioning of terrestrial ecosystems (Bardgett and Wardle 2010, Bardgett and van der Putten 2014, Wagg et al. 2014), such as water infiltration, regulation of pests and pathogens, food security, and the release of nutrients from soil organic matter (Wall et al. 2015). Energy, water, and latitude determine the alpha-diversity of above-ground organisms (plants and animals) at a global scale (Hawkins et al. 2003). However, understanding the controlling factors of soil biodiversity is still a major challenge for ecologists (Decaëns 2010, van den Hoogen et al. 2019). For instance, a recent study found that variation in earthworm diversity was mainly associated with changes in soil pH and soil carbon at small spatial scales, while precipitation and temperature were more important at a global scale (Phillips et al. 2019). Furthermore, plants and soil organisms are tightly linked through multiple direct and indirect pathways (Wardle et al. 2004). Therefore, the uncertainty of the diversity and community composition of below-ground organisms is strongly tied to the spatial scale of study due to variation in climatic, soil, and vegetation properties (Bardgett and van der Putten 2014).

Nematodes are the most abundant multi-cellular animals in soil (Bongers and Ferris 1999) and are a highly diverse group of invertebrates (Hugot et al. 2001). It is estimated that 4.4 × 10^20^ nematodes inhabit surface soils across the world, with higher abundances in sub-Arctic regions than in temperate or tropical regions, and that the amount of carbon respired by soil nematodes is equivalent to roughly 15% of carbon emissions from fossil fuel use, or around 2.2% of the total carbon emissions from soils (van den Hoogen et al. 2019). A study on the community structure of soil nematodes that included different ecosystem types did not find a good predictor of family diversity at the plot level, but nematode family richness was related to latitude at a global scale (Nielsen et al. 2014). Other studies showed that at regional scales, soil nematode community composition was mainly influenced by climatic conditions, that is, precipitation in grassland ecosystems (Chen et al. 2015, 2016; Xiong et al. 2020) and temperature in coastal ecosystems (Wu et al. 2016). Further, a recent study from forest ecosystems suggested that climate factors were the drivers of nematode community composition at the regional scale, while terrain and soil characteristics were the drivers of nematode community composition at local scales (Xiao et al. 2021). However, the study by Xiao et al. (2021) was based on a limited number of observations (i.e., sampling at three points only). So far, few studies have focused on soil nematode communities in forested ecosystems along a hydrothermal gradient. Notably, no studies have comprehensively assessed nematode diversity or community composition and their drivers in forest soils across climatic zones using standardised sampling approaches. Understanding the determinants of biodiversity in forest soils is however critical to better predict how forest ecosystems will respond to multiple environmental changes.

In the present study, we collected nematodes in forest soils from various climatic zones (i.e., temperate, warm-temperate, and tropical) to explore soil nematode distribution patterns and diversity, as well as their drivers, using boosted regression trees models (Elith et al. 2008). Boosted regression tree (BRT) models draw on insights and techniques from both statistical and machine learning methods and are a powerful alternative to other models for both explanation and prediction (Elith et al. 2008). Given that the basal resources of soil organisms are provided by plant roots, root exudates, and litter (Coleman et al. 2004), and given the slower litter decomposition rates at higher latitudes (Meentemeyer 1978) and faster root turnover rates at lower latitudes (Gill and Jackson 2000), we hypothesized that the drivers of soil nematode composition and diversity would vary across climatic zones. For instance, faster root turnover rates in tropical forest soils may provide more abundant food sources for plant-feeding nematodes. We also hypothesized that the soil nematode communities in temperate and tropical forest soils would be particularly sensitive to climatic factors. For instance, global warming may lead to an increase in nematode diversity at higher latitudes and a decrease in nematode diversity at lower latitudes because warmer temperatures would be closer to their physiological optima at higher latitudes and closer to their thermal limits at lower latitudes (Zhao et al. 2021).

## Material and Methods

### Sampling locations

A regional-scale field investigation was conducted to collect soil nematode samples from seven nature reserves, including 13 forest sites in China across a latitudinal gradient ranging from 21°36’ N to 47°11’ N, covering all major zones of forest vegetation in eastern China. From north to south, the seven nature reserves (and typical vegetation type) are: Liangshui (boreal forest), Changbaishan (mixed coniferous-broad leaf forest), Donglingshan (deciduous broad-leaved forest), Jigongshan (deciduous broad-leaved forest), Tiantongshan (evergreen broadleaved forest), Badagongshan (evergreen/deciduous broad-leaved mixed forest), and Xishuangbanna (tropical monsoon forest (Fig. 1). Generally, the northern hemisphere temperate zone refers to the region between the Tropic of Cancer (approximately 23.5° N) to the Arctic Circle (approximately 66.5°N) and the tropics is the region between the Tropic of Cancer (approximately 23.5° N) and the Tropic of Capricorn (approximately 23.5° S) (McColltoll 2005). Therefore, based on the climatic distribution of the nature reserves, we divided the gradient into three climatic zone: the forests in Liangshui (47°11’ N) and Changbaishan (42°22’ N) were classified as temperate forests, the forests in Xishuangbanna (from 21°36’ N to 21°57’ N) were classified as tropical forests, and the other forests (from 29°46’ N to 39°57’ N) were classified as warm-temperate forests.

**Fig. 1.**
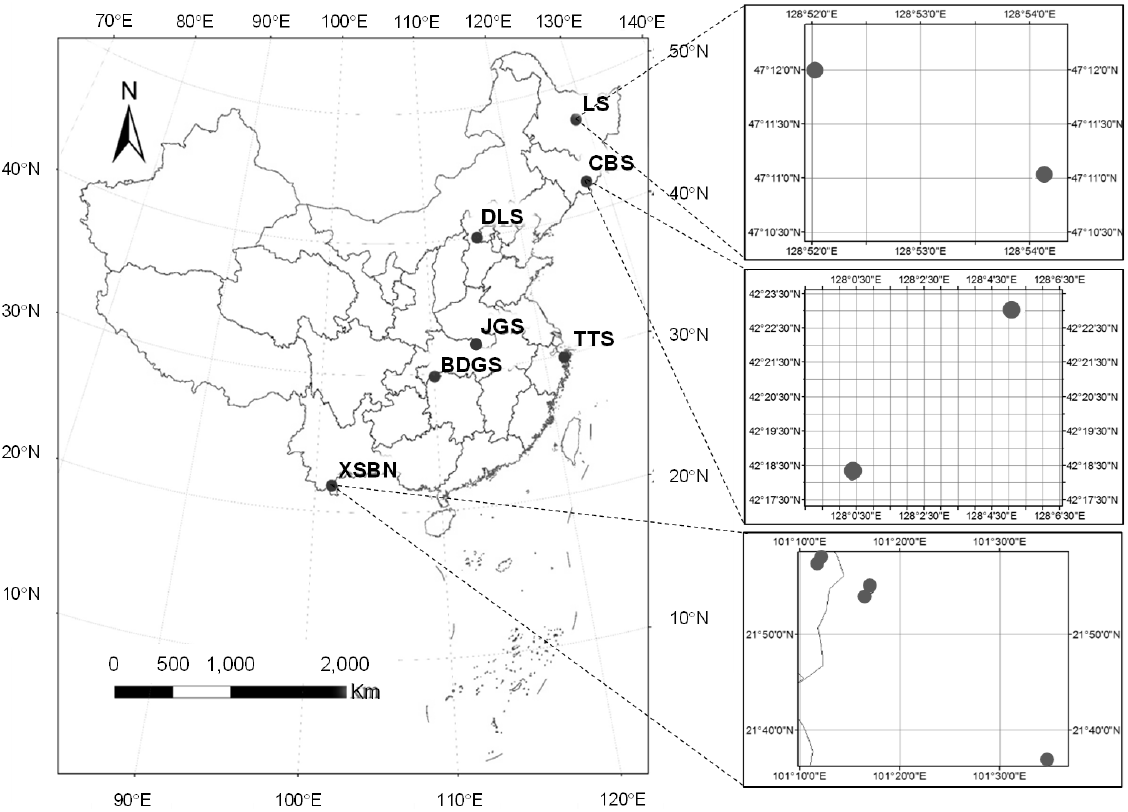
The forest sampling sites across different climatic zones in China. Dots refer to the sampling sites for field investigation, which spanned across tropical forests (Xishuangbanna, XSBN), warm-temperate forests (Donglingshan, DLS; Jigongshan, JGS; Tiantongshan, TTS; Badagongshan, BDGS), and temperate forests (Liangshui, LS; Changbaishan CBS).

### Soil sampling

In total, we selected 13 forest sites: one site in Donglingshan, Jigongshan, Tiantongshan, and Badagongshan, two sites in Liangshui and in Changbaishan, and five sites in Xishuangbanna. This resulted in four sites in temperate forest, four sites in warm-temperate forests, and five sites in tropical forest. These forest sites were selected from larger areas of forest that are representatives for the regions. The vegetation types were clearly different between any two forest sites in the same nature reserve (Appendix S1: Table S4). Three 20 × 20 m sampling plots, separated by more than 100 m, were randomly established in each forest site, resulting in a total of 39 sampling plots. Sampling was conducted between August and October 2014, during the growing season when productivity was highest. Five composite soil samples were collected from each sampling plot. Specifically, in each plot, five subplots (5 × 5 m) were established: one in each corner and one in the center. For each subplot, one composite sample comprised of five soil cores was collected from the upper 0–10 cm of soil. Litter was removed from the soil surface before soil samples were taken. Visible roots in the soil samples were picked out by hand. As a result, we collected a total of 195 samples at the subplot level (i.e., 13 sites × 3 sampling plots × 5 composite soil samples). These samples were subsequently used for analyses of physicochemical properties and soil nematodes (see below).

### Climate data

For each site, the mean annual precipitation (MAP) and mean annual temperature (MAT) for the period 1970–2000 were obtained from the WorldClim database (http://www.worldclim.org/current) (Hijmans et al. 2005).

### Plant community and productivity data

For each forest site in the tropical and temperate climatic zones, woody plant community data were obtained during one or two field surveys from 2004 to 2010; these data were provided by the nature reserves. Given the greater plant diversity and more diverse forest types in tropical forests, four 20 × 20 m plant community survey plots were randomly selected for each forest site in the tropical climatic zone while three 20 × 20 m plant community survey plots were randomly selected for each forest site in the temperate climatic zone. For the warm-temperate forests, plant community data were unfortunately unavailable.

We defined alpha-diversity of woody plant communities as the species richness of a single 20 × 20 m plot and gamma-diversity as the total richness of those plots at each forest site. Beta-diversity was defined as the species turnover among the plots at each forest site, and was measured as Beta = 1-alpha_mean_/gamma (Whittaker 1960; Kraft et al. 2011). We note that diversity indices of woody plants for warm-temperate forests could not be calculated due to the absence of data. The photosynthetic accumulation of carbon by plants per unit time, known as gross primary productivity (GPP), was estimated from satellite images of the Moderate Resolution Imaging Spectroradiometer (MODIS) according to an empirical light-use efficiency (LUE) model with the following equation (Zhang et al. 2012):

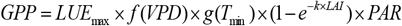

where *LUE*_max_ is the maximum light-use efficiency, *f*(*VPD*) is the scalar of daily vapor pressure deficit, *g*(*T*_min_) is the scalar of daily minimum air temperature, *e* is the natural constant, *k* is the empirical constant in calculating the probability of radiation transmission through the canopy at zenith angle, PAR is the photosynthetically active radiation absorbed by the canopy, and LAI is the leaf area index. All of these biome physical parameters are quantified based on the MODIS land cover classification system using a biome property look-up table. Given that the long-term accumulation of plant-derived carbon can also influence soil nematodes, we decided to use the dataset backwards 15 years from the date of soil sampling. Hereto, we examined the gross primary productivity from 2000–2014 for each site by using the GPP parameters from the MOD17A2H product. This global GPP product estimates the cumulative 8-day composite of values with a 500 m spatial resolution based on an empirical light-use efficiency model (Zhao et al. 2005, Zhao et al. 2010, Wang et al. 2017). In addition, to match the time of soil sampling, we selected the GPP data from August to October of each year from 2000 to 2014.

### Soil analysis

Total soil organic carbon, total soil nitrogen, total soil phosphorus, and soil pH were measured using standard methods of the Chinese Ecosystem Research Network (Liu 1996). Briefly, soil water content was measured gravimetrically by drying fresh soil at 105°C for 48 h. Soil pH was measured with a 1:2.5 (w/v) ratio of soil to deionized water using a pH meter (Mettler Toledo, Shanghai, China). Soil organic carbon was oxidized by a solution of 0.133 M K_2_Cr_2_O_7_– 18.4 M H_2_SO_4_ in an oil bath and then the excess K_2_Cr_2_O_7_ was titrated with 0.2 M FeSO_4_. Total soil nitrogen content was measured using an ultraviolet spectrophotometer after Kjeldahl digestion. Total soil phosphorus content was determined by H_2_SO_4_–HClO_4_ fusion, followed by the Mo-Sb anti-spectrophotography method.

Soil nematodes were extracted from a 30 g subsample of each composite soil sample using a sugar centrifugation method (Coleman et al. 1999) for determination of nematode abundance, biomass, and community composition. After fixation in 4% formalin solution, the nematodes were counted under an inverted microscope, and for each sample the first 100 individuals encountered were identified to the genus or family level and were classified into trophic groups: plant-feeders, bacterial feeders, fungal feeders, predators, and omnivores (Yeates et al. 1993). All nematodes were identified when the total number of nematodes was lower than 100 individuals. In this study, we defined nematode alpha-diversity as the genus or family richness in a single 5 × 5 m subplot and gamma-diversity as the total richness of the five subplots in each forest plot. Beta-diversity was defined as the turnover in taxon composition among the five subplots in each plot, and was measured as Beta = 1-alpha_mean_/gamma (Whittaker 1960, Kraft et al. 2011), where alpha_mean_ is the mean alpha diversity of each plot (N = 5).

### Statistical analysis

Given the hierarchical structure of our soil sampling protocol, linear mixed effects models (LMMs) were used to test the effects of climatic zone on soil and nematode properties, using the R-package lmerTest (Kuznetsova et al. 2017). For models testing the effects of climatic zone on soil pH, soil organic carbon, total soil nitrogen, total soil phosphorus, nematode alpha-diversity, total nematode biomass, and abundance and relative abundance of different nematode trophic groups, we fitted forest site and plot as random effects. For models testing the effects of climatic zone on nematode beta-diversity, nematode gamma-diversity, and the biomass proportion of different nematode trophic groups, we fitted forest site as a random effect. Post-hoc tests were used to compare differences among climatic zones.

For all LMMs, restricted maximum likelihood estimates of the parameters were determined using the lmer function in the lme4 package in R (Bates et al. 2015). For the LMMs, we calculated the marginal and conditional *R*^2^ values, which account for fixed and fixed plus random effects, respectively. Akaike Information Criteria (AIC) were used to test the random effects. Specifically, if there was no difference between the AIC value of the model with fixed plus random effects and the AIC value of the model with only fixed effects, then random effects were considered to be unimportant. On the other hand, if the difference in AIC value between the model with fixed plus random effects and the model with only fixed effects is large, then the random effects cannot be ignored.

WorldClim and MODIS does not allow to obtain climate and GPP differences among sub-plots within a plot and among some plots within a forest site. Therefore, one-way ANOVA was used to test the effects of climatic zone on MAP, MAT and GPP. Levene’s tests were performed to test for homogeneity of variance.

Furthermore, the data on woody plant alpha-diversity were derived from surveys at the plot level. We therefore calculated gamma-diversity and beta-diversity of woody plants at the forest site level. Thus, different from our hierarchical data on soil properties (i.e., subplots, plots, sites and climatic zones), the data on plant diversity are lacking a clear hierarchy. Hence, independent-samples t-tests were employed to compare differences in alpha-, beta-, and gamma-diversity of woody plants between the temperate zone and tropical zone. All ANOVAs and t-tests were performed using SPSS 18.0 software (SPSS Inc., Chicago, IL, USA). Statistical significance was set at *p* < 0.05.

In order to determine which variables (soil pH, soil organic carbon, total soil nitrogen, total soil phosphorus, GPP, alpha-diversity of woody plants, MAP, MAT) were most influential in driving total nematode biomass, nematode alpha-diversity, total nematode abundance and the abundance of different trophic groups, the relative importance of each variable was tested using boosted regression tree (BRT) models (Elith et al. 2008) using the gbm package in R (Ridgeway 2006). Specifically, this method combines regression trees (where models are built according to constrained clustering rules based on how recursive partitioning of a quantitative variable minimizes the within-group sums of squares under the constraining of a certain explanatory variable) and boosting algorithm (i.e., a numerical optimization technique for minimizing the loss function by stage-wisely adding a new tree). To prevent overfitting, the model used a penalized forward stepwise search and cross-validation method to identify the optimal number of decision trees (Elith et al. 2008). In BRT models, the settings of tree complexity, learning rate, bagging fraction and cross-validation folds influence the model fit (De’ath 2007; Elith et al. 2008). Therefore, multiple model processes that involved training, validation and testing were fitted with different combinations of parameter settings. Finally, models showing the smallest cross-validated relative error (CVRE) were selected as optimal models.

We ran two sets of models. First, we ran eight models (for total nematode biomass, nematode alpha-diversity, the abundance of total, plant-feeding, bacterial-feeding, fungal-feeding, omnivorous, and predatory nematodes) based on data from all individual sub-plots across all forest sites (n = 195). The optimal model settings for each dependent variable were determined by running 108 BRT processes with the following combinations of settings: tree complexities of 2, 3, 4 and 5; learning rate of 0.01, 0.005 and 0.001; bag fraction of 0.4, 0.5 and 0.6; 5-, 8- and 10-fold cross-validations. Similarly, we ran eight models (for total nematode biomass, nematode alpha-diversity, the abundance of total, plant-feeding, bacterial-feeding, fungal-feeding, omnivorous, and predatory nematodes) based on data from individual sub-plots in temperate (n = 60), warm-temperate (n = 60), and tropical (n = 75) forest sites. For these analyses, the optimal model settings for each dependent variable were determined by running 12 BRT processes with the following combinations of settings: tree complexity of 2; learning rate of 0.01, 0.005, 0.001, and 0.0001; bag fraction of 0.7; 5-, 8- and 10-fold cross-validations. In all of the above analyses, we used the same climate data for all five subplots within each sampling plot because WorldClim does not allow to detect climate differences among sub-plots at such a small spatial scale. Similarly, the MODIS GPP data have a spatial resolution of 500 m, while the area of a plot is 400 m^2^. The GPP data were obtained based on the latitude and longitude of each sampling plot. We then used the same GPP data for all five subplots within each sampling plot. In the present study, the plots for plant surveys do not exactly match with the plots for nematode sampling. Thus, for each forest site, we averaged the plant diversity values at the plot level and then used this value as a proxy of plant diversity at the subplot level.

Second, we ran eleven models (for nematode biomass, nematode alpha-, beta-, and gamma-diversity, mean alpha-diversity of nematodes at plot level, nematode richness index, and the abundance of total, plant-feeding, bacterial-feeding, fungal-feeding, omnivorous, and predatory nematodes) based on plot-level data across temperate, warm-temperate, and tropical zones (n = 39). In total, 36 BRT models for each dependent variable were fitted with the combinations of the following settings: tree complexities of 2, 3, 4 and 5; learning rate of 0.01, 0.005 and 0.001; bag fraction of 0.75; 5-, 8- and 10-fold cross-validations. The relative influence of predictor variables on the response variables was estimated based on the number of times a predictor variable was selected for splitting, weighted by the squared improvement to the model as a result of each split and averaged over all trees (De’ath 2007, Elith et al. 2008).

## Results

### Climate, plant, and soil properties

From tropical to temperate forests, mean annual precipitation (MAP) and mean annual temperature (MAT) gradually decreased, but soil organic carbon gradually increased (Table 1; Appendix S1: Tables S1, S2b). Total soil nitrogen, total soil phosphorus and nematode beta-diversity differed among climatic zones. Specifically, total soil nitrogen and total soil phosphorus were greater in temperate forests than in warm-temperate and tropical forests, while there was no difference between warm-temperate and tropical forests (Table 1; Appendix S1: Table S2c, S2d).

**Table 1.**
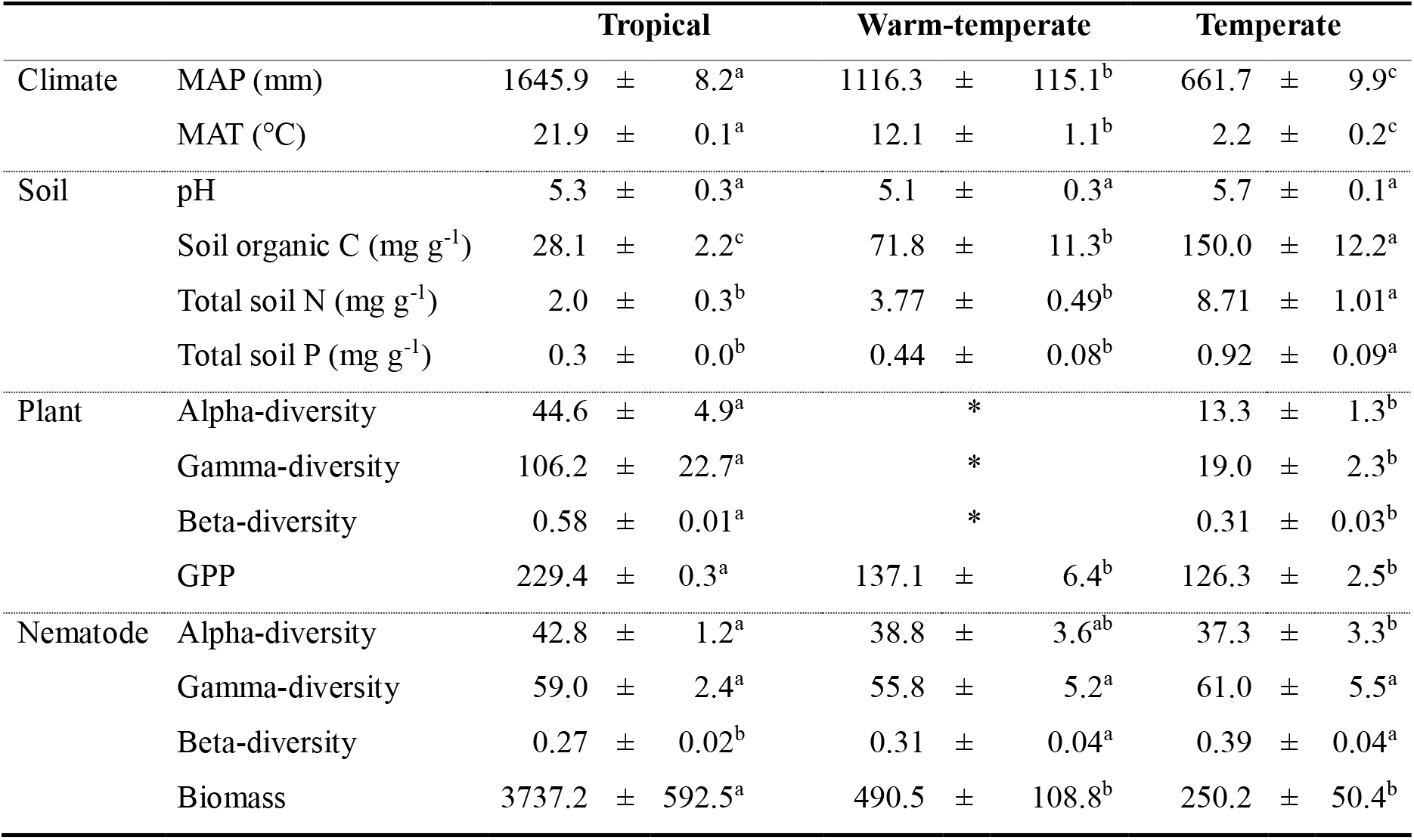
Climate, soil, plant and nematode properties for tropical, warm-temperate and temperate forest sites. Values are means ± SE for nematode beta- and gamma-diversity, and woody plant beta- and gamma diversity (n = 4 in temperate forests (4 sites), and n = 5 in tropical forests (5 sites)); for woody plant alpha-diversity (n = 12 in temperate forests (4 sites × 3 plots), and n = 20 in tropical forests (5 sites × 4 plots)); for climatic properties, soil physio-chemical properties, GPP, nematode alpha-diversity and nematode biomass (n = 12 in temperate and warm-temperate forests (4 sites × 3 plots), and n = 15 in tropical forests (5 sites × 3 plots). Different letters in a row indicate significant differences among tropical, warm-temperate, and temperate forests (post-hoc tests after one-way ANOVA for MAP, MAT and GPP, independent-samples t-test for alpha-, beta- and gamma-diversity of woody plants, and post-hoc test after linear mixed effects models for soil and nematode properties, *P* < 0.05). MAP, mean annual precipitation; MAT, mean annual temperature; *, diversity indices for warm-temperate could not be calculated due to the absence of plant data.

Nematode beta-diversity was greater in temperate and warm-temperate forests than in tropical forests, but we did not find differences in nematode beta-diversity between warm-temperate and temperate forests (Table 1; Appendix S1: Table S2g). Nematode alpha-diversity was greater in tropical forests than in temperate forests (Table 1; Appendix S1: Table S2e). Additionally, we did not find significant differences among climatic zones in nematode gamma-diversity and soil pH (Table 1; Appendix S1: Table S2f, S2a). In general, nematode biomass was greater in tropical forests than in warm-temperate and temperate forests (Table 1; Appendix S1: Table S2h). Furthermore, woody plant alpha-diversity, beta-diversity and gamma-diversity were greater in tropical than in temperate forests (Table 1; Appendix S1: Table S1).

### Drivers of nematode diversity, abundance, and biomass

At a regional scale, i.e., across tropical, warm-temperate, and temperate forests, soil organic C was more important in influencing nematode alpha-diversity, beta-diversity, gamma-diversity, and total abundance than mean annual precipitation (MAP), mean annual temperature (MAT), or vegetation properties (gross primary productivity and diversity of woody plants) (Figs. 2, 3). Mean annual temperature and total soil phosphorus in temperate forests and MAP in tropical forests were particularly important in influencing nematode alpha-diversity and trophic groups, while soil pH was the most important variable in influencing nematode alpha-diversity and trophic groups in warm-temperate forests. Interestingly, the alpha-diversity of woody plants had a greater influence on nematode alpha-diversity in tropical forests than in temperate forests. Additionally, MAP and total soil phosphorus were the most important variables in influencing nematode biomass in tropical and temperate forests, respectively (Fig. 2). Furthermore, the highest total abundance of nematodes was found in tropical forests, followed by warm-temperate forests and temperate forests (Fig. 4; Appendix S1: Table S5). Moreover, we found a greater proportional biomass of plant-feeding nematodes in tropical forests than in warm-temperate and temperate forests (Fig. 4; Appendix S1: Table S3m). We also found that warm-temperate forests were dominated by omnivorous nematodes (Fig. 4; Appendix S1: Table S3d, S3p), and that the biomass proportion and relative abundance of fungal-feeding nematodes was lower in tropical forests than in warm-temperate and temperate forests (Fig. 4; Appendix S1: Table S3h, S3n).

**Fig. 2.**
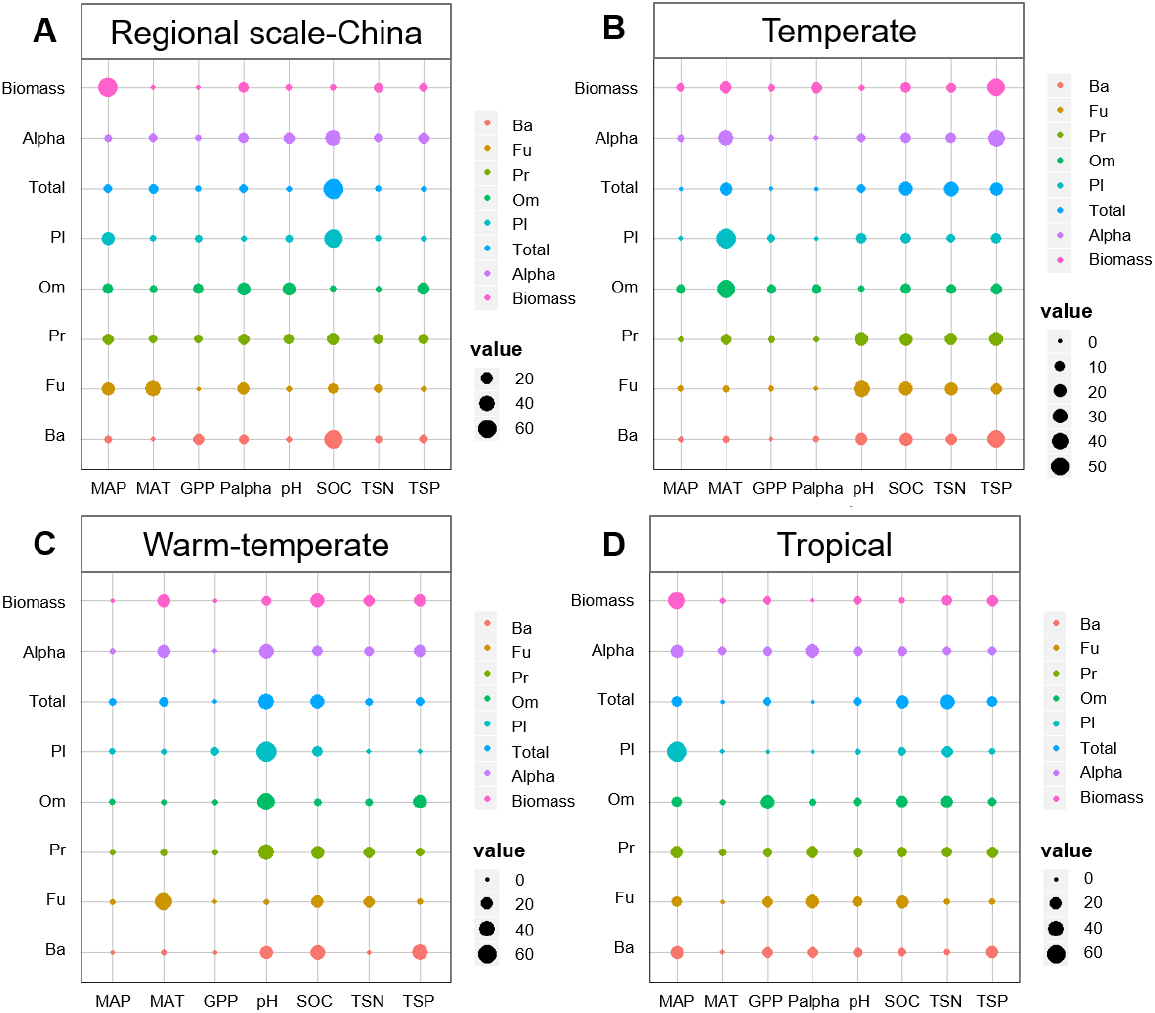
The relative importance of the seven or eight variables from the eight models based on subplot-level data at the regional scale (n = 195) (A) and separately for tropical (n = 75), warm-temperate (n = 60), and temperate (n=60) forests (B-D) in explaining soil nematode variables. Rows show the results of each model (from top to bottom, nematode biomass (Biomass), nematode alpha-diversity (Alpha), the total abundance of nematodes (Total), and the abundance of plant-feeding (Pl), omnivorous (Om), predatory (Pr), fungal-feeding (Fu), and bacterial-feeding (Ba) nematodes). Columns represent the variables that are present in the model. Within each row, the size of the circles is proportional to the relative importance of the variables. Palpha refers to the alpha-diversity of woody plants.

**Fig. 3.**
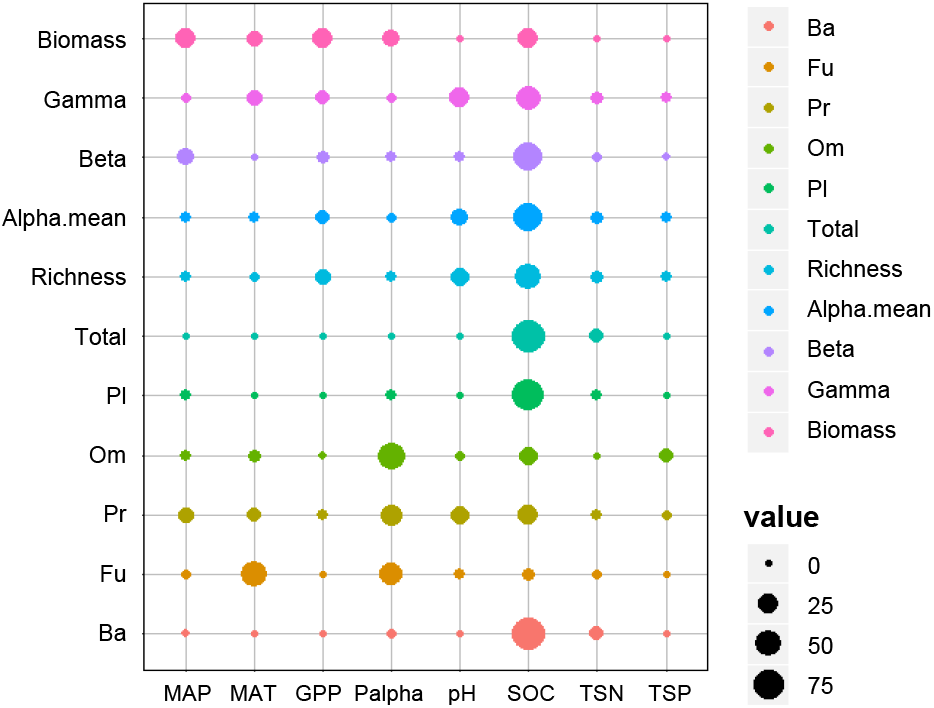
The importance of the eight variables from the eleven models based on plot-level data across temperate, warm-temperate, and tropical forest sites (n = 39) in explaining soil nematode variables. Rows show the results of each model (from top to bottom, nematode biomass (Biomass), nematode gamma-diversity (Gamma), nematode beta-diversity (Beta), nematode alpha-diversity (Alpha.mean), nematode taxon richness (Richness), the total abundance of nematodes (Total), and the abundance of plant-feeding (Pl), omnivorous (Om), predatory (Pr), fungal-feeding (Fu), and bacterial-feeding (Ba) nematodes). Columns represent the variables that are present in the model. Within each row, the size of the circles is proportional to the relative importance of the variables. Palpha refers to the alpha-diversity of woody plants.

**Fig. 4.**
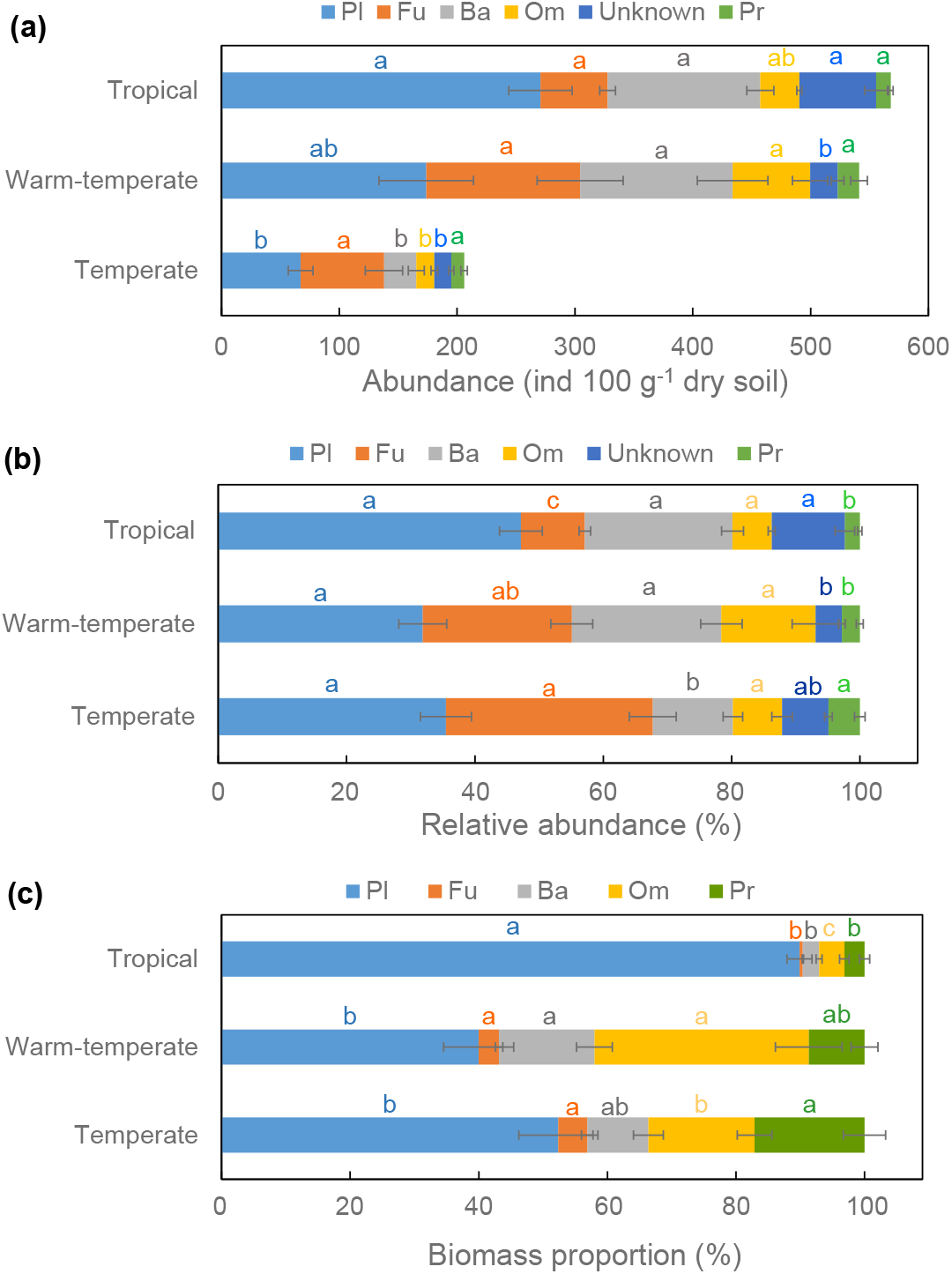
The abundance (a), relative abundance (b), and biomass proportion (c) of plant-feeding (Pl), fungal-feeding (Fu), bacterial-feeding (Ba), omnivorous (Om), unknown and predatory (Pr) nematodes for tropical, warm-temperate, and temperate forests. Notably, ‘unknown’ nematodes referred to unidentified nematodes and were not included in panel c because nematode biomass was estimated based on the identified nematodes, and biomass of unknown nematodes could not be estimated accurately. Data are means ± SE (n = 12 for temperate and warm-temperate forest plots, n = 15 for tropical forest plots). Within each panel, different lowercase letters denote significant (P < 0.05) differences among climatic zones based on post-hoc tests after linear mixed effects models.

## Discussion

In the present study, we found that the drivers of nematode diversity, biomass and abundance varied across climatic zones, and that nematode diversity, biomass and the abundance of different nematode trophic groups varied more strongly with climatic factors in temperate and tropical forests and with soil properties in warm-temperate forests. However, latitudinal variation in gross primary productivity (GPP) accumulation rate did not reflect variation in nematode abundance and diversity. Interestingly, plant diversity showed similar latitudinal patterns as the alpha-diversity of nematodes, but plant diversity did not reflect beta- and gamma-diversity of nematodes at the regional scale.

Studies on nematode diversity that focussed on marine nematode assemblages have shown that nematode diversity typically decreases with increasing latitude (Lambshead et al. 2002; Nicholas and Trueman 2005, Lee and Riveros 2012). However, patterns observed in marine environments are not easy to translate to terrestrial ecosystems. We found that nematode alpha-diversity was greater in tropical than in temperate forests, but we did not find statistical differences in nematode alpha-diversity between warm-temperate forests and tropical or temperate forests. On the contrary, we found that nematode beta-diversity was lower in tropical than in temperate and warm-temperate forests, but we did not find statistical differences in nematode beta-diversity between warm-temperate forests and temperate forests. Finally, we did not find significant differences among climatic zones in nematode gamma-diversity.

The extreme heterogeneity of soil environments at different spatial and temporal scales could promote the diversity of soil organisms (Bardgett et al. 2005). For instance, soil biodiversity at a small scale can be affected by soil compactness (Decaëns 2010), and small-scale soil spatial heterogeneity has been shown to promote nematode diversity (Nielsen et al. 2010). On the other hand, at a spatial larger scale, soil fertility may largely affect the composition of soil communities (Laliberté et al. 2017). In the present study, soil organic C increased from tropical to temperate forests. Furthermore, soil properties may be influenced in part by the structure and composition of the current vegetation through root and litter input (Coleman et al. 2004), but at the same time soil formation is a long-term process and the development of soil properties is partly independent of the current vegetation and may also reflect past environmental conditions (Wall et al. 2005). This could explain why we found that soil properties exerted a greater impact on soil nematode abundance and diversity than aboveground plant GPP and diversity across climatic zones.

On the other hand, plant-feeding nematodes are likely to be strongly affected by the current plant productivity (and biomass) and community composition as they directly feed on plant roots or root hairs (Ferris and Bongers 2006, Neher 2010). However, in our dataset, plant productivity did not explain much of the variation in the abundance of plant-feeding nematodes. We did find however that the GPP and the biomass of nematodes in tropical forests were greater than those in warm-temperate and temperate forests. Interestingly, the alpha-diversity of woody plants had a greater influence on nematode alpha-diversity in tropical forests than in temperate forests. Also, the biomass of plant-feeding nematodes in tropical forests accounted for more than 90% of the total nematode biomass, and it was greater than the biomass of plant-feeding nematodes in warm temperate and temperate forests. This indicates that the plant-feeding nematodes and their diversity, relative to other nematode trophic groups, are more likely to be controlled by the bottom-up effects of plant productivity and plant diversity (Ferris and Bongers 2006, Neher 2010). Here, plant roots might represent a particularly important food resource for nematodes because root turnover rates are greater at low latitudes than at high latitudes (Gill and Jackson 2000). Furthermore, the diversity in root morphological traits may be greatest at lower latitudes, where plant diversity is highest (Ma et al. 2018); this may cause greater heterogeneity in soil environments and potentially stimulate greater nematode diversity in tropical forests.

Availability of food and water resources, coupled with a suitable temperature, constitute the principal factors that control nematode abundance and distribution (Yeates 2004). The normal temperature at which nematodes are active is roughly between 5 and 30 °C with the optimal activity temperature being about 20 °C (Wharton 2004). In our study, the variation in mean monthly temperature ranged from -20 to 20 °C in temperate forests, which is greater than in warm-temperate forests (−8 to 26 °C) and tropical forests (about 16 to 26 °C) (Appendix S1: Table S6), and this may be one of the reasons why MAT explained more of the variation in nematode alpha-diversity and in the abundance of different nematode trophic groups in temperate forests. Also, at higher latitudes, litter decomposition rate is slower (Meentemeyer 1978) and accumulated litter may create variable microclimatic conditions for soil fauna. Meanwhile, the greater temperature fluctuations in temperate forests may result in more variability in resource availability in time and space. As a result, greater litter accumulation and temperature fluctuations in temperate forests may promote heterogeneity in nematodes assemblages, which could explain the increase in nematode beta-diversity with increasing latitude.

In addition, the movement, development and survival of nematodes in soil are also regulated by soil porosity and water potential (Barbercheck and Duncan 2004). For instance, variation in precipitation can alter the composition of soil nematode communities (Franco et al. 2019), which may result in changes in nematode diversity. Reduced aeration in saturated soil is often detrimental to nematodes and it has been shown that the developmental rate and movement of many nematodes is highest when soil water content changes from saturated to slightly drier states (Barbercheck and Duncan 2004). In our studied system, there is greater variation in mean monthly precipitation in tropical forests (ranging from 16 to 315 mm) relative to warm-temperate forests (2 to 231 mm) and temperate forests (3 to 170 mm) (Appendix S1: Table S7), and this may explain why MAP in tropical forests explains more of the variability in soil nematode alpha-diversity and in the abundance of different nematode trophic groups compared to the other forest types.

## Conclusions

Given that nematodes are the most abundant animals on Earth and play key roles in regulating carbon and nutrient dynamics, and reflecting biological activity in soils (van den Hoogen et al. 2019), understanding the drivers of nematode diversity in forest soils is essential to better predict how forest ecosystems respond to multiple environmental changes. Based on a large-scale investigation of soil nematode diversity and the abundance of different nematode trophic groups and their drivers in forested ecosystems, we conclude that across a hydrothermal gradient from temperate to tropical climates, nematode abundance and diversity were mainly influenced by soil organic carbon. Within climatic zones, however, mean annual temperature and total soil phosphorus in temperate, soil pH in warm temperate, and mean annual precipitation in tropic were more important drivers. This demonstrates that the drivers of soil nematode diversity in forested ecosystems are affected by the spatial scale and climatic conditions considered.

In temperate forests the magnitude of variation in temperature may be an important driver of nematode diversity, while in tropical forests the magnitude of variation in precipitation may be more important. This indicates a latitudinal shift in the main climatic variables controlling soil nematode communities. Our data further indicated that the alpha- and beta-diversity of woody plant and nematode alpha-diversity were greater in the tropical forests than in the temperate forests, while nematode beta-diversity was lower in the tropical forests than in the temperate forests. This demonstrates a complex and scale-dependent relationship between plant diversity and soil biodiversity. These findings also have consequences for the scales of observation of area–diversity relationships (Decaëns 2010). For those forests with greater nematode beta-diversity, such as temperate forests, increasing the area of sampling may enhance the probability of detecting rare species. Therefore, for better understanding the drivers of soil biodiversity distribution, it is crucial to explore the area–diversity relationships of soil organisms across various ecosystems.

## Supporting information

Appendix S1 for Drivers of nematode diversity in forest soils across climatic zones

## Data Availability

All data are presented in the paper.

## Acknowledgments

We acknowledge all members from the Institute of Geographic Sciences and Natural Resources Research, Chinese Academy of Sciences, and Peking University for their help with the fieldwork. This study was sponsored by the Natural Science Foundation of China (31971534, U1904204, U1804101) and the Innovation Scientists and Technicians Troop Construction Projects of Henan Province (grant number 182101510005). We are also very grateful for the support from the nature reserves.

## Author Contributions

All authors contributed intellectual input and assistance to this study. D.G. and S.F. designed the study. Y.S. wrote the first draft with help of P.K. Sample collection, and soil chemical and nematode analysis were carried out by C.M., Z.W., T.L., and Y.S. Climate and plant data were collected by Y.H., C.M. and Y.S. Statistical analyses were carried out by Y.S., Y.C. and Z.W. Assistance in data interpretation was provided by Y.S., P.K., Y.H., Y.C., C.Z. and W.Z.

## References

Barbercheck, M. E., and L. Duncan. 2004. Abiotic Factors. Pages 309–343 in Gaugler R., and A. L. Bilgrami, editors. Nematode Behaviour. CABI Publishing, Wallingford, Oxfordshire. UK.

Bardgett, R. D. and W. H. van der Putten 2014. Belowground biodiversity and ecosystem functioning. Nature 515: 505–511.

Bardgett, R. D., and D. A. Wardle. 2010. Aboveground-Belowground Linkages: Biotic Interactions, Ecosystem Processes, and Global Change. Oxford University Press. Oxford. UK.

Bardgett, R. D., G. W. Yeates, and J. M. Anderson. 2005. Patterns and determinants of soil biological diversity. Pages 100–118 in R. D. Bardgett, M. B. Usher, and D. W. Hopkins, editors. Biological Diversity and Function in Soils. Cambridge University Press, Cambridge. UK.

Bates, D., M. Mächler, B. Bolker, and S. Walker. 2015. “Fitting Linear Mixed-Effects Models Using lme4. “ Journal of Statistical Software 67: 1–48. doi: 10.18637/jss.v067.i01.

Bongers, T. and H. Ferris. 1999. Nematode community structure as a bioindicator in environmental monitoring. Trends in Ecology & Evolution 14: 224–228.

Chen, D., J. Cheng, P. Chu, J. Mi, and Y. Bai. 2016. Effect of diversity on biomass across grasslands on the mongolian plateau: contrasting effects between plants and soil nematodes. Journal of Biogeography 43: 955–966.

Chen, D., J. Cheng, P. Chu, S. Hu, Y. Xie, I. Tuvshintogtokh, and Y. Bai. 2015. Regional-scale patterns of soil microbes and nematodes across grasslands on the Mongolian plateau: relationships with climate, soil, and plants. Ecography 38: 622–631.

Coleman, D.C., J. M. Blair, E. T. Elliott, and D. H. Wall. 1999. Soil invertebrates. Pages 349–377 in G. P. Robertson, D. C. Coleman, C. S. Bedsoe, and P. Sollins, editors. Standard Soil Methods for Long-term Ecological Research. Oxford University Press. New York. USA.

Coleman, D.C., D. A. Crossley Jr., and P. F. Hendrix. 2004. Fundamentals of Soil Ecology. Second edition. Elsevier Academic Press, Burlington, San Diego, London.

De ‘ath, G. 2007. Boosted trees for ecological modeling and prediction, Ecology 88, 243–251.

Decaëns, T. 2010. Macroecological patterns in soil communities. Global Ecology and Biogeography 19: 287–302.

Elith, J., J. R. Leathwick, and T. Hastie. 2008. A working guide to boosted regression trees. Journal of Animal Ecology 77: 802–813.

Ferris, H., and T. Bongers. 2006. Nematode indicators of organic enrichment. Journal of Nematology 38: 3–12.

Franco, A.L.C., L. A. Gherardi,C. M. D. Tomasel,W. S. Andriuzzi, and D. H. Wall. 2019. Drought suppresses soil predators and promotes root herbivores in mesic, but not in xeric grasslands. Proceedings of the National Academy of Sciences USA 116: 12883–12888. doi: 10.1073/pnas.1900572116.

Gill, R. A., and R. B. Jackson. 2000. Global patterns of root turnover for terrestrial ecosystems. New Phytologist 147: 13–31.

Hawkins, B.A. et al. 2003. Energy, water, and broad-scale geographic patterns of species richness. Ecology 84: 3105–3117.

Hijmans, R. J., S. E. Cameron, J. L. Parra, P. G. Jones, and A. Jarvis. 2005. Very high resolution interpolated climate surfaces for global land areas. International Journal of Climatology 25: 1965–1978.

Hugot, J. P., P. Baujard, and S. Morand. 2001. Biodiversity in helminths and nematodes as a field of study: an overview. Nematology 3: 199–208.

Kraft, N. J. et al. 2011. Disentangling the drivers of β diversity along latitudinal and elevational gradients. Science 333: 1755–1758.

Kuznetsova, A., P. B. Brockhoff, and R. H. B. Christensen. 2017. “lmerTest Package: Tests in Linear Mixed Effects Models. “ Journal of Statistical Software 82: 1–26. doi: 10.18637/jss.v082.i13.

Laliberté, E., P. Kardol., R. K. Didham, F. P. Teste, B. L. Turner, and D. A. Wardle. 2017. Soil fertility shapes belowground food webs across a regional climate gradient. Ecology Letters 20: 1273–1284.

Lambshead, P., C. J. Brown, T. J. Ferrero, N. J. Mitchell, C. R. Smith, L. E. Hawkins, and J. Tietjen. 2002. Latitudinal diversity patterns of deep-sea marine nematodes and organic fluxes: A test from the central equatorial Pacific. Marine Ecology Progress Series 236: 129–135.

Lee, M. R., and M. Riveros. 2012. Latitudinal trends in the species richness of free-living marine nematode assemblages from exposed sandy beaches along the coast of Chile (18– 42°S). Marine Ecology 33: 317–325.

Liu, G. 1996. Soil Physical and Chemical Analysis & Description of Soil Profiles. China Standard, Beijing, China (in Chinese).

Ma, Z.Q. et al. 2018. Evolutionary history resolves global organization of root functional traits. Nature 555: 94–97.

McColltoll, R. W. 2005. Encyclopedia of World Geography, Volume 1. Facts on File Library of World Geography. Facts on File, New York, USA.

Meentemeyer, V. 1978. Macroclimate and lignin control of litter decomposition rates. Ecology 59, 465–472.

Neher, D. A. 2010. Ecology of plant and free-living nematodes in natural and agricultural soil. Annual Review of Phytopathology 48: 371–394.

Nicholas, W. L., and J. Trueman. 2005. Biodiversity of marine nematodes in Australian sandy beaches from tropical and temperate regions. Biodiversity and Conservation 14: 823–839.

Nielsen, U.N., E. Ayres, D. H. Wall, G. Li, R. D. Bardgett, T. Wu, and J. R. Garey. 2014. Global-scale patterns of assemblage structure of soil nematodes in relation to climate and ecosystem properties. Global Ecology and Biogeography 23: 968–978.

Nielsen, U.N., G. H. R. Osler,C. D. Campbell,R. Neilson,D. F. R. P. Burslem, and R. vander Wal. 2010. The enigma of soil animal species diversity revisited: The role of small-scale heterogeneity. PLoS One 5: e11567.

Phillips, H.R.P. et al. 2019. Global distribution of earthworm diversity. Science 366: 480–485.

Ridgeway, G. 2006. Generalized boosted regression models. Documentation on the R Package ‘gbm ‘, version 1· 5–7.

van den Hoogen, J. et al. 2019. Soil nematode abundance and functional group composition at a global scale. Nature 572: 194–198.

Wagg, C., S. F. Bender, F. Widmer, and M. G. A. van der Heijden. 2014. Soil biodiversity and soil community composition determine ecosystem multifunctionality. Proceedings of the National Academy of Sciences USA 111: 5266–5270.

Wall, D. H., A. H. Fitter, and E. A. Paul. 2005. Developing new perspectives from advances in soil biodiversity research. Pages 3–27 in R. D. Bardgett, M. B. Usher, and D.W. Hopkins, editors. Biological Diversity and Function in Soils. Cambridge University Press, Cambridge, UK.

Wall, D.H., U.N. Nielsen, and J. Six. 2015. Soil biodiversity and human health. Nature 528: 69–76.

Wang, L, H. Zhu, A. Lin, L. Zou, W. Qin, and Q. Du. 2017. Evaluation of the latest MODIS GPP products across multiple biomes using global eddy covariance flux data. Remote Sensing 9: 418; doi:10.3390/rs9050418.

Wardle, D. A., R. D. Bardgett, J. N. Klironomos, H. Setälä, W. H. van der Putten, and D. H. Wall. 2004. Ecological linkages between aboveground and belowground biota. Science 304: 1629–1633.

Wharton, D.A. 2004. Survival Strategies. Pages 371–399 in R. Gaugler, and A. L. Bilgrami, editors. Nematode Behaviour. CABI Publishing, Wallingford, Oxfordshire. UK.

Whittaker, R.H. 1960. Vegetation of the Siskiyou Mountains, Oregon and California. Ecological Monographs 30: 279–338.

Wu, J., H. Chen, and Y. Zhang 2016. Latitudinal variation in nematode diversity and ecological roles along the Chinese coast. Ecology and Evolution 6: 8018–8027.

Xiao, H., W. Wang, S. Xia, Z. Li, J. Gan, and X. Yang. 2021. Distributional patterns of soil nematodes in relation to environmental variables in forest ecosystems. Soil Ecology Letters 3: 115–124.

Xiong, D. et al. 2020. Nonlinear responses of soil nematode community composition to increasing aridity. Global Ecology and Biogeography 29: 117–126.

Yeates, G.W. 2004. Ecological and Behavioural Adaptations. Pages 1–24 in R. Gaugler, and A. L. Bilgrami, editors. Nematode Behaviour. CABI Publishing, Wallingford, Oxfordshire. UK.

Yeates, G.W., T. Bongers, R. G. M. De Goede, D. W. Freckman, and S. S. Georgieva. 1993. Feeding habits in soil nematode families and genera—an outline for soil ecologists. Journal of Nematology 25: 315–331.

Zhang, F, J. M. Chen, J. Chen, C. M. Gough, T. A. Martin, and D. Dragoni. 2012. Evaluating spatial and temporal patterns of MODIS GPP over the conterminous U.S. against flux measurements and a process model. Remote Sensing of Environment 124: 717–729.

Zhao, M, F. A. Heinsch, R. R. Nemani, and S.W. Running. 2005. Improvements of the MODIS terrestrial gross and net primary production global data set. Remote Sensing of Environment 95: 164–176.

Zhao, M, and S.W. Running. 2010. Drought-induced reduction in global terrestrial net primary production from 2000 through 2009. Science 329: 940–943.

Zhao, L. et al. 2021. The effects of plant resource inputs on the energy flux of soil nematodes are affected by climate and plant resource type. Soil Ecology Letters 3: 134–144.

