## Appendix S1 for Drivers of nematode diversity in forest soils across climatic zones for "Drivers of nematode diversity in forest soils across climatic zones"

**Drivers of nematode diversity in forest soils across climatic zones**

Yuanhu Shao, Zuyan Wang, Tao Liu, Paul Kardol, Chengen Ma, Yonghong Hu, Yang Cui, Cancan Zhao, Weixin Zhang, Dali Guo, Shenglei Fu

**This document includes:
Table S1.** Results (*F*-values and *P*-values) of one-way ANOVAs or t-tests (*t*-values and *P*-values) testing the effects of climatic zone on climate (MAP and MAT) and plant (alpha-, beta- and gamma-diversity of woody plants and GPP) properties.

**Table S2.** Results of linear mixed models testing the effects of climatic zone on soil pH, soil organic carbon, total soil nitrogen, total soil phosphorus (with forest site as random effect, n = 13; and with plot as random effect, n = 39) and nematode alpha-diversity and total nematode biomass (with forest site as random effect, n = 13; and with plot as random effect, n = 39), nematode beta- and gamma-diversity (with forest site as random effect, n = 13).

**Table S3.** Results of linear mixed models testing the effects of climatic zone on abundance, relative abundance (with forest site as random effect, n = 13; and with plot as random effect, n = 39) and the biomass proportion of different nematode trophic groups (with forest site as random effect, n = 13).

**Table S4.** Composition of dominant tree species for forest sites in temperate, warm-temperate, and tropical climatic zones.

**Table S5.** The abundance (individuals/100 g dry soil) of nematode genera/families for tropical, warm-temperate, and temperate forest sites.

**Table S6.** The mean monthly temperature (°C) for the 39 sampling plots in temperate, warm-temperate, and tropical climatic zones.

**Table S7.** The mean monthly precipitation (mm) for the 39 sampling plots in temperate, warm-temperate, and tropical climatic zones.

**Table S1.** Results (*F*-values and *P*-values) of one-way ANOVAs or t-tests (*t*-values and *P*-values) testing the effects of climatic zone on climate (MAP and MAT) and plant (alpha-, beta- and gamma-diversity of woody plants and GPP) properties.

| **Variable** | **Results of one-way ANOVA or t-test** | |
| --- | --- | --- |
| MAP (mm) | *F*_(2,36)_ = 66.2 | *P* < 0.001 |
| MAT (°C) | *F*_(2,36)_ = 275.5 | *P* < 0.001 |
| Alpha-diversity | *t*_(30)_ = 4.88 | *P* < 0.001 |
| Gamma-diversity | *t*_(7)_ = 3.38 | *P* = 0.01 |
| Beta-diversity | *t*_(7)_ = 9.11 | *P* < 0.001 |
| GPP (g C m^-2^ month^-1^) | *F*_(2,36)_ = 255.7 | *P* < 0.001 |

**Table S2.** Results of linear mixed models testing the effects of climatic zone on soil pH, soil organic carbon, total soil nitrogen, total soil phosphorus (with forest site as random effect, n = 13; and with plot as random effect, n = 39) and nematode alpha-diversity and total nematode biomass (with forest site as random effect, n = 13; and with plot as random effect, n = 39), nematode beta- and gamma-diversity (with forest site as random effect, n = 13). The marginal R^2^ indicates the variance explained by the fixed effects, while the conditional R^2^ indicates the variance explained by both fixed and random effects. The AIC value of the model with random effects are shown.

| a. **Soil pH** | | | | |
| --- | --- | --- | --- | --- |
| Fixed effects | Estimate | SE | t | *p*-value |
| Warm-temperate vs. temperate | -0.5957 | 0.6889 | -0.865 | 0.407 |
| Tropical vs. temperate | -0.3667 | 0.6535 | -0.561 | 0.587 |
| Tropical vs. warm-temperate | 0.2289 | 0.6535 | 0.35 | 0.733 |
| Random effects variance | Plot | Forest site | Residual |  |
|  | 0.03968 | 0.93087 | 0.07611 |  |
| Model fit | Marginal R^2^ | Conditional R^2^ | AIC |  |
|  | 0.051 | 0.931 | 160.53 |  |
| b. **Soil organic carbon** | | | | |
| Fixed effects | Estimate | SE | t | *p*-value |
| Warm-temperate vs. temperate | -78.24 | 17.7 | -4.42 | 0.00129 ** |
| Tropical vs. temperate | -121.88 | 16.8 | -7.257 | 2.74e-05 *** |
| Tropical vs. warm-temperate | -43.64 | 16.8 | -2.599 | 0.026555 * |
| Random effects variance | Plot | Forest site | Residual |  |
|  | 553.3 | 386.1 | 843.1 |  |
| Model fit | Marginal R^2^ | Conditional R^2^ | AIC |  |
|  | 0.591 | 0.807 | 1924.8 |  |
| c. **Total soil nitrogen** | | | | |
| Fixed effects | Estimate | SE | t | *p*-value |
| Warm-temperate vs. temperate | -4.933 | 1.419 | -3.476 | 0.005962 ** |
| Tropical vs. temperate | -6.689 | 1.346 | -4.968 | 0.000564 *** |
| Tropical vs. warm temperate | -1.756 | 1.346 | -1.304 | 0.2215 |
| Random effects variance | Plot | Forest site | Residual |  |
|  | 2.028 | 3.22 | 1.998 |  |
| Model fit | Marginal R^2^ | Conditional R^2^ | AIC |  |
|  | 0.525 | 0.869 | 783.26 |  |
| d. **Total soil phosphorus** | | | | |
| Fixed effects | Estimate | SE | t | *p*-value |
| Warm-temperate vs. temperate | -0.4814 | 0.1537 | -3.133 | 0.01063 * |
| Tropical vs. temperate | -0.6237 | 0.1458 | -4.278 | 0.00162 ** |
| Tropical vs. warm-temperate | -0.1422 | 0.1458 | -0.976 | 0.35219 |
| Random effects variance | Plot | Forest site | Residual |  |
|  | 0.028 | 0.037 | 0.014 |  |
| Model fit | Marginal R^2^ | Conditional R^2^ | AIC |  |
|  | 0.47 | 0.904 | -142.823 |  |
| e. **Nematode alpha-diversity** | | | | |
| Fixed effects | Estimate | SE | t | *p*-value |
| Warm-temperate vs. temperate | 3.388 | 3.596 | 0.942 | 0.3682 |
| Tropical vs. temperate | 9.221 | 3.412 | 2.703 | 0.0221 * |
| Tropical vs. warm-temperate | 5.833 | 3.408 | 1.711 | 0.118 |
| Random effects variance | Plot | Forest site | Residual |  |
|  | 8.654 | 20.992 | 28.513 |  |
| Model fit | Marginal R^2^ | Conditional R^2^ | AIC |  |
|  | 0.207 | 0.611 | 1231 |  |
| f. **Nematode gamma-diversity** | | | | |
| Fixed effects | Estimate | SE | t | *p*-value |
| Warm-temperate vs. temperate | 1.583 | 5.647 | 0.280 | 0.785 |
| Tropical vs. temperate | 5.550 | 5.357 | 1.036 | 0.325 |
| Tropical vs. warm-temperate | 3.967 | 5.357 | 0.740 | 0.476 |
| Random effects variance | Forest site | Residual |  |  |
|  | 43.11 | 61.97 |  |  |
| Model fit | Marginal R^2^ | Conditional R^2^ | AIC |  |
|  | 0.053 | 0.442 | 279.68 |  |
| g. **Nematode beta-diversity** | | | | |
| Fixed effects | Estimate | SE | t | *p*-value |
| Warm-temperate vs. temperate | -0.05733 | 0.03024 | -1.896 | 0.087 |
| Tropical vs. temperate | -0.16062 | 0.02869 | -5.598 | 0.000228 *** |
| Tropical vs. warm-temperate | -0.10328 | 0.02869 | -3.600 | 0.00485 ** |
| Random effects variance | Forest site | Residual |  |  |
|  | 0.0008995 | 0.0027889 |  |  |
| Model fit | Marginal R^2^ | Conditional R^2^ | AIC |  |
|  | 0.563 | 0.669 | -85.146 |  |
| h. **Nematode biomass** | | | | |
| Fixed effects | Estimate | SE | t | *p*-value |
| Warm-temperate vs. temperate | 246.219 | 933.346 | 0.264 | 0.7973 |
| Tropical vs. temperate | 3562.81 | 885.516 | 4.023 | 0.00243 ** |
| Tropical vs. warm temperate | 3316.591 | 883.55 | 3.754 | 0.00382 ** |
| Random effects variance | Plot | Forest site | Residual |  |
|  | 75655 | 1382735 | 4824231 |  |
| Model fit | Marginal R^2^ | Conditional R^2^ | AIC |  |
|  | 0.311 | 0.471 | 3451.4 |  |

**Table S3.** Results of linear mixed models testing the effects of climatic zone on abundance, relative abundance (with forest site as random effect, n = 13; and with plot as random effect, n = 39) and the biomass proportion of different nematode trophic groups (with forest site as random effect, n = 13). The Marginal R^2^ indicates the variance explained by the fixed effects, while the Conditional R^2^ indicates the variance explained by both fixed and random effects. The AIC value of the model with random effects are shown.

| a. **Plant-feeding nematode abundance** | | | | |
| --- | --- | --- | --- | --- |
| Fixed effects | Estimate | SE | t | *p*-value |
| Warm-temperate vs. temperate | 121.3 | 74.22 | 1.634 | 0.1331 |
| Tropical vs. temperate | 203.75 | 70.41 | 2.894 | 0.0159 * |
| Tropical vs. warm-temperate | 82.46 | 70.31 | 1.173 | 0.26809 |
| Random effects variance | Plot | Forest site | Residual |  |
|  | 0 | 9669 | 19451 |  |
| Model fit | Marginal R^2^ | Conditional R^2^ | AIC |  |
|  | 0.196 | 0.463 | 2423 |  |
| b. **Fungal-feeding nematode abundance** | | | | |
| Fixed effects | Estimate | SE | t | *p*-value |
| Warm-temperate vs. temperate | 62.56 | 43 | 1.455 | 0.1762 |
| Tropical vs. temperate | -13.1 | 40.79 | -0.321 | 0.7547 |
| Tropical vs. warm-temperate | -75.656 | 40.746 | -1.857 | 0.0930 . |
| Random effects variance | Plot | Forest site | Residual |  |
|  | 3552 | 2206 | 4409 |  |
| Model fit | Marginal R^2^ | Conditional R^2^ | AIC |  |
|  | 0.096 | 0.608 | 2190.7 |  |
| c. **Bacterial-feeding nematode abundance** | | | | |
| Fixed effects | Estimate | SE | t | *p*-value |
| Warm-temperate vs. temperate | 103.12 | 33.52 | 3.077 | 0.01166 * |
| Tropical vs. temperate | 102.31 | 31.8 | 3.218 | 0.00917 ** |
| Tropical vs. warm-temperate | -0.8008 | 31.7279 | -0.025 | 0.980362 |
| Random effects variance | Plot | Forest site | Residual |  |
|  | 2262 | 1114 | 5437 |  |
| Model fit | Marginal R^2^ | Conditional R^2^ | AIC |  |
|  | 0.202 | 0.507 | 2210.2 |  |
| d. **Omnivorous nematode abundance** | | | | |
| Fixed effects | Estimate | SE | t | *p*-value |
| Warm-temperate vs. temperate | 46.86 | 16.04 | 2.921 | 0.0152 * |
| Tropical vs. temperate | 15.76 | 15.22 | 1.036 | 0.3245 |
| Tropical vs. warm-temperate | -31.09 | 15.19 | -2.047 | 0.067838 . |
| Random effects variance | Plot | Forest site | Residual |  |
|  | 439 | 289.6 | 1132.7 |  |
| Model fit | Marginal R^2^ | Conditional R^2^ | AIC |  |
|  | 0.159 | 0.488 | 1916.6 |  |
| e. **Unknown nematode abundance** | | | | |
| Fixed effects | Estimate | SE | t | *p*-value |
| Warm-temperate vs. temperate | 7.821 | 15.409 | 0.508 | 0.62271 |
| Tropical vs. temperate | 55.655 | 14.619 | 3.807 | 0.00342 ** |
| Tropical vs. warm-temperate | 47.834 | 14.606 | 3.275 | 0.00835 ** |
| Random effects variance | Plot | Forest site | Residual |  |
|  | 73.92 | 418.1 | 462.37 |  |
| Model fit | Marginal R^2^ | Conditional R^2^ | AIC |  |
|  | 0.404 | 0.711 | 1744.6 |  |
| f. **Predatory nematode abundance** | | | | |
| Fixed effects | Estimate | SE | t | *p*-value |
| Warm-temperate vs. temperate | -1.872 | 3.738 | -0.501 | 0.62728 |
| Tropical vs. temperate | 2.228 | 3.546 | 0.628 | 0.54383 |
| Tropical vs. warm-temperate | 4.1 | 3.536 | 1.16 | 0.2733 |
| Random effects variance | Plot | Forest site | Residual |  |
|  | 21.12 | 14.18 | 96.66 |  |
| Model fit | Marginal R^2^ | Conditional R^2^ | AIC |  |
|  | 0.022 | 0.284 | 1443.1 |  |
| g. **Relative abundance of plant-feeding nematodes** | | | | |
| Fixed effects | Estimate | SE | t | *p*-value |
| Warm-temperate vs. temperate | -0.02031 | 0.07931 | -0.256 | 0.803 |
| Tropical vs. temperate | 0.09619 | 0.07524 | 1.278 | 0.23 |
| Tropical vs. warm-temperate | 0.1165 | 0.07516 | 1.55 | 0.152186 |
| Random effects variance | Plot | Forest site | Residual |  |
|  | 0.0053 | 0.0098 | 0.0144 |  |
| Model fit | Marginal R^2^ | Conditional R^2^ | AIC |  |
|  | 0.086 | 0.554 | -186.1 |  |
| h. **Relative abundance of fungal-feeding nematodes** | | | | |
| Fixed effects | Estimate | SE | t | *p*-value |
| Warm-temperate vs. temperate | -0.0819 | 0.06034 | -1.357 | 0.20433 |
| Tropical vs. temperate | -0.22522 | 0.05725 | -3.934 | 0.00276 ** |
| Tropical vs. warm-temperate | -0.14332 | 0.05718 | -2.507 | 0.031037 * |
| Random effects variance | Plot | Forest site | Residual |  |
|  | 0.0055 | 0.0048 | 0.0095 |  |
| Model fit | Marginal R^2^ | Conditional R^2^ | AIC |  |
|  | 0.314 | 0.67 | -255.95 |  |
| i. **Relative abundance of bacterial-feeding nematodes** | | | | |
| Fixed effects | Estimate | SE | t | *p*-value |
| Warm-temperate vs. temperate | 0.09669 | 0.03129 | 3.091 | 0.003814 ** |
| Tropical vs. temperate | 0.11406 | 0.02968 | 3.843 | 0.000469 *** |
| Tropical vs. warm-temperate | 0.01737 | 0.02961 | 0.587 | 0.56102 |
| Random effects variance | Plot | Forest site | Residual |  |
|  | 0.0047 | 0 | 0.0056 |  |
| Model fit | Marginal R^2^ | Conditional R^2^ | AIC |  |
|  | 0.191 | 0.56 | -355.56 |  |
| j. **Relative abundance of omnivorous nematodes** | | | | |
| Fixed effects | Estimate | SE | t | *p*-value |
| Warm-temperate vs. temperate | 0.06496 | 0.036802 | 1.765 | 0.108 |
| Tropical vs. temperate | -0.009687 | 0.034915 | -0.277 | 0.787 |
| Tropical vs. warm-temperate | -0.07465 | 0.03488 | -2.14 | 0.058166 . |
| Random effects variance | Plot | Forest site | Residual |  |
|  | 0.003 | 0.0015 | 0.003 |  |
| Model fit | Marginal R^2^ | Conditional R^2^ | AIC |  |
|  | 0.125 | 0.65 | -458.05 |  |
| k. **Relative abundance of unknown nematodes** | | | | |
| Fixed effects | Estimate | SE | t | *p*-value |
| Warm-temperate vs. temperate | -0.03038 | 0.02343 | -1.297 | 0.22352 |
| Tropical vs. temperate | 0.04773 | 0.02223 | 2.147 | 0.05704 . |
| Tropical vs. warm-temperate | 0.07811 | 0.02219 | 3.521 | 0.0055 ** |
| Random effects variance | Plot | Forest site | Residual |  |
|  | 0.0002 | 0.0009 | 0.0025 |  |
| Model fit | Marginal R^2^ | Conditional R^2^ | AIC |  |
|  | 0.238 | 0.466 | -542.18 |  |
| l. **Relative abundance of predatory nematodes** | | | | |
| Fixed effects | Estimate | SE | t | *p*-value |
| Warm-temperate vs. temperate | -0.029551 | 0.00849 | -3.481 | 0.00583 ** |
| Tropical vs. temperate | -0.023148 | 0.008056 | -2.874 | 0.01640 * |
| Tropical vs. warm-temperate | 0.006403 | 0.008022 | 0.798 | 0.44338 |
| Random effects variance | Plot | Forest site | Residual |  |
|  | 0.0001 | 0.000056 | 0.0007 |  |
| Model fit | Marginal R^2^ | Conditional R^2^ | AIC |  |
|  | 0.144 | 0.31 | -773.3 |  |
| m. **Biomass proportion of plant-feeding nematodes** | | | | |
| Fixed effects | Estimate | SE | t | *p*-value |
| Warm-temperate vs. temperate | -12.381 | 7.904 | -1.566 | 0.148331 |
| Tropical vs. temperate | 37.523 | 7.499 | 5.004 | 0.000534 *** |
| Tropical vs. warm-temperate | 49.904 | 7.499 | 6.655 | 0.000057 *** |
| Random effects variance | Forest site | Residual |  |  |
|  | 48.74 | 228.65 |  |  |
| Model fit | Marginal R^2^ | Conditional R^2^ | AIC |  |
|  | 0.638 | 0.701 | 320.34 |  |
| n. **Biomass proportion of fungal-feeding nematodes** | | | | |
| Fixed effects | Estimate | SE | t | *p*-value |
| Warm-temperate vs. temperate | -1.3275 | 1.1853 | -1.120 | 0.288911 |
| Tropical vs. temperate | -4.0028 | 1.1245 | -3.560 | 0.005184 ** |
| Tropical vs. warm-temperate | -2.6753 | 1.1245 | -2.379 | 0.0387 * |
| Random effects variance | Forest site | Residual |  |  |
|  | 1.875 | 2.805 |  |  |
| Model fit | Marginal R^2^ | Conditional R^2^ | AIC |  |
|  | 0.39 | 0.634 | 167.98 |  |
| o. **Biomass proportion of bacterial-feeding nematodes** | | | | |
| Fixed effects | Estimate | SE | t | *p*-value |
| Warm-temperate vs. temperate | 5.343 | 3.431 | 1.557 | 0.15046 |
| Tropical vs. temperate | -6.954 | 3.255 | -2.137 | 0.05837 . |
| Tropical vs. warm-temperate | -12.296 | 3.255 | -3.778 | 0.003613 ** |
| Random effects variance | Forest site | Residual |  |  |
|  | 9.726 | 41.438 |  |  |
| Model fit | Marginal R^2^ | Conditional R^2^ | AIC |  |
|  | 0.345 | 0.47 | 259.24 |  |
| p. **Biomass proportion of omnivorous nematodes** | | | | |
| Fixed effects | Estimate | SE | t | *p*-value |
| Warm-temperate vs. temperate | 16.802 | 4.611 | 3.644 | 0.00084 *** |
| Tropical vs. temperate | -12.601 | 4.374 | -2.881 | 0.00664 ** |
| Tropical vs. warm-temperate | -29.403 | 4.374 | -6.722 | 0.000000076*** |
| Random effects variance | Forest site | Residual |  |  |
|  | 0 | 127.6 |  |  |
| Model fit | Marginal R^2^ | Conditional R^2^ | AIC |  |
|  | 0.543 | 0.543 | 294.39 |  |
| q. **Biomass proportion of predatory nematodes** | | | | |
| Fixed effects | Estimate | SE | t | *p*-value |
| Warm-temperate vs. temperate | -8.436 | 3.942 | -2.140 | 0.05804 . |
| Tropical vs. temperate | -13.962 | 3.740 | -3.733 | 0.00389 ** |
| Tropical vs. warm-temperate | -5.526 | 3.740 | -1.478 | 0.170 |
| Random effects variance | Forest site | Residual |  |  |
|  | 15.87 | 45.61 |  |  |
| Model fit | Marginal R^2^ | Conditional R^2^ | AIC |  |
|  | 0.358 | 0.524 | 264.52 |  |

**Table S4.** Composition of dominant tree species for forest sites in temperate, warm-temperate, and tropical climatic zones.

| **Climatic zone** | **Nature reserve** | **Forest type** | **Dominant tree species** |
| --- | --- | --- | --- |
| Temperate forests | Liangshui | Spruce-fir forest | *Picea asperata*  *Abies fabri* |
|  |  | Mixed broad-leaved and Korean pine forest | *Acer pictum*  *Pinus koraiensis*  *Acer tegmentosum* |
|  | Changbaishan | Mixed coniferous-broad leaf forest | *Acer barbinerve*  *Acer pseudosieboldianum*  *Pinus koraiensis*  *Tilia amurensis* |
|  |  | Secondary *Populus davidiana*-  *Betula platyphylla* forest | *Betula platyphylla*  *Tilia amurensis*  *Populus daviana* |
| Warm-temperate forests | Donglingshan | Deciduous broad-leaved forest | *Quercus wutaishansea*  *Acer mono*  *Betula duburica* |
|  | Jigongshan | Deciduous broad-leaved forest | *Quercus acutissima*  *Quercus variabilis*  *Liquidambar formosana* |
|  | Badagongshan | Evergreen/deciduous broad-leaved mixed forest | *Cyclobalanopsis multinervis*  *Fagus lucida*  *Cyclobalanopsis gracilis*  *Litsea, elongate* |
|  | Tiantongshan | Evergreen broadleaved forest | *Lithocarpus harlandii*  *Schima superba*  *Litsea elongate*  *Machilus leptophylla* |
| Tropical forests | Xishuangbanna | Tropical monsoon forest over limestone | *Cleistanthus sumatranus*  *Celtis philippensis* |
|  |  | Tropical monsoon evergreen broad­-leaved forest | *Castanopsis echidnocarpa*  *Syzygium oblatum*  *Millettia leptobotrya*  *Aporusa yunnanensis* |
|  |  | Tropical secondary forest | *Syzygium oblatum,*  *Millettia leptobotrya*  *Phoebe lanceolata*  *Castanopsis echidnocarpa* |
|  |  | Tropical secondary monsoon forest | *Barringtonia fusicarpa*  *Millettia leptobotrya*  *Mezzettiopsis creaghii*  *Chisocheton paniculatus* |
|  |  | Tropical monsoon forest | *Parashorea chinensis*  *Phoebe lanceolata,*  *Garcinia cowa*  *Baccaurea ramiflora* |

**Table S5.** The abundance (individuals/100 g dry soil) of nematode genera/families for tropical, warm-temperate, and temperate forest sites. Values are means ± SE.

| Trophic group | Genus/Family | **Tropical** | **Warm-temperate** | **Temperate** |
| --- | --- | --- | --- | --- |
| Bacterial-feeding | *Achromadora* | 0.00 ± 0.00 | 0.00 ±0.00 | 0.07 ± 0.07 |
|  | *Acrobeles* | 0.00 ± 0.00 | 16.56 ± 12.18 | 0.07 ± 0.07 |
|  | *Acrobeloides* | 13.95 ± 2.24 | 3.33 ± 0.98 | 1.63 ± 0.57 |
|  | *Alaimus* | 38.50 ± 4.46 | 28.23 ± 12.20 | 3.61 ± 1.84 |
|  | *Anaplectus* | 0.00 ± 0.00 | 0.05 ± 0.04 | 0.51 ± 0.22 |
|  | *Aphanolaimus* | 7.29 ± 1.62 | 5.01 ± 3.70 | 0.10 ± 0.10 |
|  | *Bastiania* | 0.00 ± 0.00 | 4.53 ± 2.03 | 2.82 ± 1.19 |
|  | *Brevibucca* | 10.48 ± 1.33 | 0.00 ± 0.00 | 0.00 ± 0.00 |
|  | *Bunonema* | 0.00 ± 0.00 | 0.10 ± 0.09 | 0.00 ± 0.00 |
|  | Cephalobidae | 0.00 ± 0.00 | 0.05 ± 0.04 | 0.10 ± 0.10 |
|  | *Cephalobus* | 0.00 ± 0.00 | 6.93 ± 2.98 | 0.51 ± 0.17 |
|  | *Cervidellus* | 0.15 ± 0.05 | 2.70 ± 2.08 | 0.27 ± 0.12 |
|  | *Chiloplacus* | 0.10 ± 0.09 | 0.00 ± 0.00 | 0.00 ± 0.00 |
|  | *Chronogaster* | 0.00 ± 0.00 | 0.69 ± 0.38 | 0.10 ± 0.06 |
|  | *Desmolaimus* | 0.00 ± 0.00 | 0.00 ± 0.00 | 0.16 ± 0.16 |
|  | *Desmoscolex* | 1.39 ± 0.64 | 0.00 ± 0.00 | 0.00 ± 0.00 |
|  | *Eucephalobus* | 12.04 ± 4.51 | 2.24 ± 0.73 | 0.10 ± 0.10 |
|  | *Eumonhystera* | 12.09 ± 2.25 | 0.00 ± 0.00 | 0.15 ± 0.15 |
|  | *Euteratocephalus* | 0.04 ± 0.04 | 0.00 ± 0.00 | 0.05 ± 0.05 |
|  | *Heterocephalobus* | 1.10 ± 0.25 | 4.77 ± 1.23 | 0.88 ± 0.58 |
|  | *Ironus* | 0.10 ± 0.05 | 0.00 ± 0.00 | 0.05 ± 0.05 |
|  | *Leptolaimus* | 0.00 ± 0.00 | 0.00 ± 0.00 | 0.08 ± 0.08 |
|  | *Metateratocephalus* | 0.33 ± 0.14 | 0.00 ± 0.00 | 1.47 ± 0.86 |
|  | *Microlaimus* | 1.01 ± 0.33 | 0.00 ± 0.00 | 0.00 ± 0.00 |
|  | *Monhystera* | 12.24 ± 3.68 | 0.00 ± 0.00 | 0.15 ± 0.15 |
|  | *Panagrellus* | 0.64 ± 0.35 | 0.16 ± 0.14 | 0.00 ± 0.00 |
|  | *Panagrolaimus* | 0.04 ± 0.04 | 0.00 ± 0.00 | 0.33 ± 0.33 |
|  | *Paramphidelus* | 0.00 ± 0.00 | 0.08 ± 0.07 | 0.00 ± 0.00 |
|  | *Paraplectonema* | 0.00 ± 0.00 | 0.66 ± 0.45 | 0.05 ± 0.05 |
|  | *Pellioditis* | 0.44 ± 0.24 | 0.00 ± 0.00 | 0.00 ± 0.00 |
|  | *Plectonchus* | 0.00 ± 0.00 | 0.06 ± 0.05 | 0.00 ± 0.00 |
|  | *Plectus* | 1.19 ± 0.48 | 6.25 ± 4.17 | 7.31 ± 4.23 |
|  | *Prismatolaimus* | 6.61 ± 1.49 | 35.32 ± 11.91 | 3.33 ± 0.63 |
|  | *Pristionchus* | 0.05 ± 0.04 | 0.00 ± 0.00 | 0.00 ± 0.00 |
|  | Rhabditidae | 0.39 ± 0.30 | 0.00 ± 0.00 | 0.00 ± 0.00 |
|  | *Rhabditis* | 0.20 ± 0.18 | 0.55 ± 0.18 | 0.19 ± 0.13 |
|  | *Rhabdolaimus* | 5.86 ± 1.25 | 0.15 ± 0.13 | 0.10 ± 0.10 |
|  | *Stenochulus* | 0.44 ± 0.24 | 0.00 ± 0.00 | 0.00 ± 0.00 |
|  | *Teratocephalus* | 1.66 ± 1.11 | 3.21 ± 2.02 | 2.84 ± 0.96 |
|  | *Tylopharynx* | 0.05 ± 0.04 | 0.00 ± 0.00 | 0.00 ± 0.00 |
|  | *Wilsonema* | 0.56 ± 0.20 | 3.21 ± 2.63 | 0.30 ± 0.23 |
| Fungal-feeding | *Anomyctus* | 0.00 ± 0.00 | 0.08 ± 0.07 | 0.00 ± 0.00 |
|  | *Aphelenchoides* | 10.83 ± 2.05 | 2.78 ± 1.14 | 3.80 ± 1.89 |
|  | *Aphelenchus* | 0.00 ± 0.00 | 1.46 ± 1.04 | 0.12 ± 0.07 |
|  | *Bursaphelenchus* | 0.15 ± 0.13 | 0.00 ± 0.00 | 0.00 ± 0.00 |
|  | *Campydora* | 0.22 ± 0.13 | 0.05 ± 0.04 | 0.00 ± 0.00 |
|  | *Diphtherophora* | 0.27 ± 0.11 | 0.70 ± 0.61 | 0.15 ± 0.09 |
|  | *Ditylenchus* | 0.00 ± 0.00 | 2.04 ± 1.02 | 1.55 ± 0.80 |
|  | *Filenchus* | 31.12 ± 7.92 | 130.80 ± 53.48 | 68.21 ± 21.08 |
|  | *Fungiotonchium* | 0.00 ± 0.00 | 0.00 ± 0.00 | 0.05 ± 0.05 |
|  | *Nothotylenchus* | 0.00 ± 0.00 | 0.00 ± 0.00 | 0.05 ± 0.05 |
|  | *Paraphelenchus* | 0.00 ± 0.00 | 0.00 ± 0.00 | 0.05 ± 0.05 |
|  | *Seinura* | 0.09 ± 0.05 | 0.55 ± 0.48 | 0.05 ± 0.05 |
|  | *Tylencholaimellus* | 0.05 ± 0.04 | 0.00 ± 0.00 | 0.08 ± 0.08 |
|  | *Tylencholaimus* | 13.98 ± 3.02 | 0.00 ± 0.00 | 0.05 ± 0.05 |
|  | *Tylolaimophorus* | 0.00 ± 0.00 | 0.05 ± 0.04 | 0.00 ± 0.00 |
| Omnivorous | *Allodorylaimus* | 0.00 ± 0.00 | 0.00 ± 0.00 | 0.08 ± 0.08 |
|  | *Aporcelaimellus* | 1.23 ± 0.51 | 0.47 ± 0.14 | 0.27 ± 0.18 |
|  | *Belondirella* | 3.00 ± 0.56 | 0.00 ± 0.00 | 0.00 ± 0.00 |
|  | *Chrysonemoides* | 0.00 ± 0.00 | 0.50 ± 0.32 | 0.07 ± 0.07 |
|  | *Dorydorella* | 0.05 ± 0.04 | 0.00 ± 0.00 | 0.00 ± 0.00 |
|  | Dorylaimidae | 0.00 ± 0.00 | 0.48 ± 0.24 | 0.40 ± 0.21 |
|  | *Dorylaimoides* | 0.00 ± 0.00 | 0.00 ± 0.00 | 0.05 ± 0.05 |
|  | *Enchodelus* | 4.28 ± 1.10 | 0.00 ± 0.00 | 0.00 ± 0.00 |
|  | *Epidorylaimus* | 0.91 ± 0.59 | 12.68 ± 7.48 | 2.69 ± 0.82 |
|  | *Eudorylaimus* | 10.20 ± 3.03 | 12.42 ± 5.65 | 1.45 ± 0.47 |
|  | *Laimydorus* | 0.00 ± 0.00 | 0.00 ± 0.00 | 0.10 ± 0.10 |
|  | Nordiidae | 0.05 ± 0.04 | 0.00 ± 0.00 | 0.00 ± 0.00 |
|  | *Mesodorylaimus* | 5.53 ± 1.54 | 0.75 ± 0.45 | 0.66 ± 0.37 |
|  | *Microdorylaimus* | 2.93 ± 0.76 | 0.11 ± 0.06 | 0.82 ± 0.42 |
|  | *Nygolaimus* | 0.00 ± 0.00 | 0.16 ± 0.14 | 0.00 ± 0.00 |
|  | *Opisthodorylaimus* | 0.00 ± 0.00 | 0.16 ± 0.14 | 0.00 ± 0.00 |
|  | *Prodorylaimus* | 2.63 ± 1.50 | 12.29 ± 5.79 | 2.68 ± 0.68 |
|  | Qudsianematidae | 0.04 ± 0.04 | 0.00 ± 0.00 | 0.05 ± 0.05 |
|  | *Thonus* | 0.00 ± 0.00 | 5.65 ± 1.33 | 0.90 ± 0.32 |
|  | *Thornia* | 0.05 ± 0.04 | 17.03 ± 7.40 | 4.19 ± 1.75 |
|  | *Torumanawa* | 0.00 ± 0.00 | 0.00 ± 0.00 | 0.08 ± 0.08 |
| Plant-feeding | *Aglenchus* | 0.17 ± 0.11 | 2.59 ± 1.42 | 0.19 ± 0.19 |
|  | *Aorolaimus* | 0.00 ± 0.00 | 1.15 ± 0.63 | 3.25 ± 2.73 |
|  | Criconematidae | 0.00 ± 0.00 | 0.00 ± 0.00 | 0.14 ± 0.14 |
|  | *Basiria* | 3.42 ± 2.65 | 0.09 ± 0.08 | 0.05 ± 0.05 |
|  | *Boleodorus* | 1.63 ± 0.65 | 5.50 ± 2.14 | 1.13 ± 0.67 |
|  | *Cephalenchus* | 0.10 ± 0.05 | 0.05 ± 0.04 | 0.73 ± 0.62 |
|  | *Coslenchus* | 100.55 ± 22.95 | 1.58 ± 0.94 | 5.91 ± 0.93 |
|  | *Criconema* | 46.43 ± 18.02 | 0.64 ± 0.30 | 0.53 ± 0.08 |
|  | *Criconemella* | 0.34 ± 0.31 | 0.00 ± 0.00 | 0.79 ± 0.79 |
|  | *Criconemoides* | 0.88 ± 0.41 | 0.32 ± 0.27 | 0.00 ± 0.00 |
|  | *Crossonema* | 1.44 ± 0.50 | 0.33 ± 0.20 | 0.23 ± 0.14 |
|  | Heteroderidae | 0.00 ± 0.00 | 0.16 ± 0.14 | 2.15 ± 1.05 |
|  | *Deladenus* | 0.00 ± 0.00 | 0.00 ± 0.00 | 0.54 ± 0.47 |
|  | *Dolichorhynchus* | 0.00 ± 0.00 | 1.21 ± 1.05 | 0.66 ± 0.66 |
|  | *Helicotylenchus* | 6.12 ± 3.51 | 17.66 ± 9.13 | 1.64 ± 0.43 |
|  | *Hemicriconemoides* | 0.00 ± 0.00 | 0.00 ± 0.00 | 0.05 ± 0.05 |
|  | *Hemicycliophora* | 0.64 ± 0.24 | 0.00 ± 0.00 | 0.24 ± 0.14 |
|  | *Heterodera* | 0.00 ± 0.00 | 2.31 ± 1.80 | 0.00 ± 0.00 |
|  | *Hexatylus* | 0.00 ± 0.00 | 9.48 ± 3.08 | 2.56 ± 0.91 |
|  | *Histotylenchus* | 0.09 ± 0.08 | 0.00 ± 0.00 | 0.00 ± 0.00 |
|  | Hoplolaimidae | 0.00 ± 0.00 | 0.00 ± 0.00 | 0.10 ± 0.10 |
|  | *Hoplolaimus* | 0.00 ± 0.00 | 0.00 ± 0.00 | 0.15 ± 0.15 |
|  | *Lelenchus* | 4.04 ± 1.72 | 16.74 ± 11.70 | 2.73 ± 0.68 |
|  | *Longidorella* | 0.29 ± 0.21 | 2.97 ± 1.77 | 0.47 ± 0.25 |
|  | *Longidorus* | 2.86 ± 0.47 | 0.87 ± 0.40 | 0.54 ± 0.22 |
|  | *Loofia* | 0.00 ± 0.00 | 0.27 ± 0.14 | 0.00 ± 0.00 |
|  | Criconematidae | 4.59 ± 2.60 | 33.86 ± 7.84 | 5.11 ± 1.21 |
|  | *Malenchus* | 0.60 ± 0.49 | 5.33 ± 2.82 | 1.48 ± 0.66 |
|  | *Meloidogyne* | 67.33 ± 15.95 | 0.00 ± 0.00 | 0.00 ± 0.00 |
|  | *Merlinius* | 0.00 ± 0.00 | 0.92 ± 0.42 | 1.70 ± 1.08 |
|  | *Nagelus* | 0.00 ± 0.00 | 0.16 ± 0.14 | 0.08 ± 0.08 |
|  | *Neothada* | 0.00 ± 0.00 | 0.00 ± 0.00 | 0.05 ± 0.05 |
|  | *Ogma* | 1.93 ± 0.35 | 5.48 ± 1.10 | 2.18 ± 0.54 |
|  | *Pararotylenchus* | 0.00 ± 0.00 | 0.00 ± 0.00 | 0.08 ± 0.08 |
|  | *Paratrichodorus* | 0.34 ± 0.16 | 8.39 ± 4.33 | 2.09 ± 1.70 |
|  | *Paratrophurus* | 0.00 ± 0.00 | 0.75 ± 0.66 | 0.08 ± 0.08 |
|  | *Paratylenchus* | 9.64 ± 2.74 | 5.66 ± 3.42 | 1.64 ± 0.46 |
|  | *Pratylenchus* | 8.32 ± 3.55 | 1.05 ± 0.33 | 0.91 ± 0.20 |
|  | *Psilenchus* | 0.05 ± 0.04 | 0.00 ± 0.00 | 0.00 ± 0.00 |
|  | *Pungentus* | 0.95 ± 0.76 | 10.37 ± 5.09 | 0.22 ± 0.14 |
|  | *Rotylenchus* | 0.59 ± 0.53 | 45.00 ± 35.94 | 10.83 ± 5.46 |
|  | *Scutellonema* | 0.00 ± 0.00 | 0.00 ± 0.00 | 0.30 ± 0.23 |
|  | *Telotylenchus* | 0.00 ± 0.00 | 0.00 ± 0.00 | 0.24 ± 0.24 |
|  | Ecphyadophoridae | 2.02 ± 0.72 | 0.00 ± 0.00 | 0.00 ± 0.00 |
|  | *Trichodorus* | 0.71 ± 0.49 | 0.74 ± 0.42 | 0.72 ± 0.37 |
|  | Tylenchulidae | 0.23 ± 0.13 | 0.00 ± 0.00 | 0.00 ± 0.00 |
|  | *Trophurus* | 0.89 ± 0.79 | 0.00 ± 0.00 | 0.00 ± 0.00 |
|  | Tylenchidae | 4.92 ± 2.34 | 0.06 ± 0.05 | 0.05 ± 0.05 |
|  | *Tylenchorhynchus* | 0.00 ± 0.00 | 0.41 ± 0.12 | 0.87 ± 0.80 |
|  | *Tylenchus* | 0.43 ± 0.23 | 0.10 ± 0.09 | 10.24 ± 2.91 |
|  | Tylodoridae | 0.00 ± 0.00 | 0.86 ± 0.50 | 0.52 ± 0.30 |
|  | *Xiphinema* | 1.00 ± 0.61 | 0.00 ± 0.00 | 0.00 ± 0.00 |
|  | *Zygotylenchus* | 0.00 ± 0.00 | 0.00 ± 0.00 | 0.05 ± 0.05 |
| Predatory | *Afractionolaimus* | 0.05 ± 0.04 | 0.00 ± 0.00 | 0.00 ± 0.00 |
|  | *Carcharolaimus* | 0.10 ± 0.09 | 0.00 ± 0.00 | 0.00 ± 0.00 |
|  | *Clarkus* | 0.18 ± 0.10 | 1.07 ± 0.68 | 0.39 ± 0.27 |
|  | Nygolaimidae | 0.76 ± 0.32 | 0.00 ± 0.00 | 0.00 ± 0.00 |
|  | *Coomansus* | 0.42 ± 0.17 | 0.00 ± 0.00 | 0.00 ± 0.00 |
|  | *Discolaimus* | 0.00 ± 0.00 | 0.09 ± 0.08 | 0.00 ± 0.00 |
|  | *Iotonchulus* | 0.24 ± 0.22 | 0.00 ± 0.00 | 0.00 ± 0.00 |
|  | *Iotonchus* | 3.37 ± 1.16 | 0.00 ± 0.00 | 0.00 ± 0.00 |
|  | *Mononchus* | 0.05 ± 0.04 | 0.82 ± 0.30 | 0.10 ± 0.10 |
|  | *Mylonchulus* | 1.05 ± 0.47 | 0.55 ± 0.48 | 0.93 ± 0.56 |
|  | *Paractinolaimus* | 0.22 ± 0.09 | 0.47 ± 0.41 | 0.00 ± 0.00 |
|  | Mononchidae | 0.00 ± 0.00 | 0.56 ± 0.33 | 1.79 ± 0.64 |
|  | *Tobrilus* | 2.74 ± 0.74 | 0.13 ± 0.07 | 0.33 ± 0.23 |
|  | *Tripyla* | 3.77 ± 1.38 | 1.54 ± 0.96 | 3.11 ± 0.90 |
|  | *Trischistoma* | 0.00 ± 0.00 | 3.42 ± 1.44 | 4.47 ± 2.07 |
| Total (identified) |  | 503.00 ± 35.15 | 517. 81 ± 153.91 | 191.39 ± 24.62 |
| Unknown |  | 65.14 ± 10.84 | 23.40 ± 6.08 | 14.45 ± 2.29 |

**Table S6.** The mean monthly temperature (°C) for the 39 sampling plots in temperate, warm-temperate, and tropical climatic zones.

| Climatic zone | Forest site | Plot | Jan | Feb | Mar | Apr | May | Jun | Jul | Aug | Sep | Oct | Nov | Dec |
| --- | --- | --- | --- | --- | --- | --- | --- | --- | --- | --- | --- | --- | --- | --- |
| Temperate | Liangshui1 | P1 | -20.2 | -15.6 | -6.7 | 3.6 | 10.7 | 16.9 | 20.2 | 18.4 | 11.4 | 2.5 | -8.3 | -17.0 |
| Temperate | Liangshui1 | P2 | -20.2 | -15.6 | -6.7 | 3.6 | 10.7 | 16.9 | 20.2 | 18.4 | 11.4 | 2.5 | -8.3 | -17.0 |
| Temperate | Liangshui1 | P3 | -20.2 | -15.6 | -6.7 | 3.6 | 10.7 | 16.9 | 20.2 | 18.4 | 11.4 | 2.5 | -8.3 | -17.0 |
| Temperate | Liangshui2 | P1 | -20.0 | -15.4 | -6.5 | 3.8 | 10.9 | 17.2 | 20.5 | 18.8 | 11.7 | 2.7 | -8.2 | -16.9 |
| Temperate | Liangshui2 | P2 | -20.0 | -15.4 | -6.5 | 3.8 | 10.9 | 17.2 | 20.5 | 18.8 | 11.7 | 2.7 | -8.2 | -16.9 |
| Temperate | Liangshui2 | P3 | -20.0 | -15.4 | -6.6 | 3.7 | 10.8 | 17.1 | 20.4 | 18.7 | 11.6 | 2.6 | -8.2 | -16.8 |
| Temperate | Changbaishan1 | P1 | -15.3 | -12.0 | -4.7 | 4.6 | 10.9 | 15.9 | 19.6 | 18.9 | 11.8 | 5.0 | -3.7 | -11.9 |
| Temperate | Changbaishan1 | P2 | -15.4 | -12.0 | -4.7 | 4.6 | 10.9 | 15.9 | 19.7 | 18.9 | 11.8 | 5.0 | -3.7 | -11.9 |
| Temperate | Changbaishan1 | P3 | -15.4 | -12.0 | -4.7 | 4.6 | 10.9 | 15.9 | 19.7 | 18.9 | 11.8 | 5.0 | -3.7 | -11.9 |
| Temperate | Changbaishan2 | P1 | -15.4 | -12.2 | -5.5 | 3.6 | 10.3 | 15.4 | 19.1 | 18.2 | 11.3 | 4.1 | -4.1 | -11.8 |
| Temperate | Changbaishan2 | P2 | -15.4 | -12.2 | -5.5 | 3.6 | 10.3 | 15.4 | 19.0 | 18.2 | 11.3 | 4.1 | -4.0 | -11.8 |
| Temperate | Changbaishan2 | P3 | -15.4 | -12.2 | -5.5 | 3.6 | 10.3 | 15.4 | 19.0 | 18.2 | 11.3 | 4.1 | -4.0 | -11.8 |
| Warm-temperate | Donglingshan | P1 | -8.7 | -6.9 | -1.0 | 7.5 | 14.5 | 19.1 | 20.9 | 18.9 | 13.3 | 7.5 | 0.0 | -6.5 |
| Warm-temperate | Donglingshan | P2 | -8.7 | -6.9 | -1.0 | 7.5 | 14.5 | 19.1 | 20.9 | 18.9 | 13.3 | 7.5 | 0.0 | -6.5 |
| Warm-temperate | Donglingshan | P3 | -8.7 | -6.9 | -1.0 | 7.5 | 14.5 | 19.1 | 20.9 | 18.9 | 13.3 | 7.5 | 0.0 | -6.5 |
| Warm-temperate | Jigongshan | P1 | 2.5 | 4.6 | 9.3 | 15.9 | 20.9 | 25.0 | 27.5 | 26.5 | 21.7 | 16.4 | 10.3 | 4.8 |
| Warm-temperate | Jigongshan | P2 | 2.3 | 4.4 | 9.1 | 15.7 | 20.7 | 24.7 | 27.1 | 26.2 | 21.4 | 16.2 | 10.1 | 4.7 |
| Warm-temperate | Jigongshan | P3 | 2.5 | 4.6 | 9.3 | 15.9 | 20.9 | 25.0 | 27.5 | 26.5 | 21.7 | 16.4 | 10.3 | 4.8 |
| Warm-temperate | Badagongshan | P1 | 0.6 | 2.3 | 5.9 | 11.5 | 15.2 | 18.3 | 20.6 | 20.3 | 16.7 | 12.1 | 7.4 | 3.1 |
| Warm-temperate | Badagongshan | P2 | 0.6 | 2.3 | 5.9 | 11.5 | 15.2 | 18.3 | 20.6 | 20.3 | 16.7 | 12.1 | 7.4 | 3.1 |
| Warm-temperate | Badagongshan | P3 | 0.6 | 2.3 | 5.9 | 11.5 | 15.2 | 18.3 | 20.6 | 20.3 | 16.7 | 12.1 | 7.4 | 3.1 |
| Warm-temperate | Tiantongshan | P1 | 4.0 | 5.1 | 8.5 | 13.9 | 18.6 | 22.3 | 26.3 | 26.2 | 22.6 | 17.5 | 12.2 | 6.6 |
| Warm-temperate | Tiantongshan | P2 | 4.0 | 5.1 | 8.5 | 13.9 | 18.6 | 22.3 | 26.3 | 26.2 | 22.6 | 17.5 | 12.2 | 6.6 |
| Warm-temperate | Tiantongshan | P3 | 4.0 | 5.1 | 8.5 | 13.9 | 18.6 | 22.3 | 26.3 | 26.2 | 22.6 | 17.5 | 12.2 | 6.6 |
| Tropical | Xishuangbanna1 | P1 | 17.0 | 18.6 | 21.4 | 23.9 | 25.7 | 26.2 | 25.6 | 25.7 | 25.1 | 23.4 | 20.2 | 17.0 |
| Tropical | Xishuangbanna1 | P2 | 17.0 | 18.6 | 21.4 | 23.9 | 25.7 | 26.2 | 25.6 | 25.7 | 25.1 | 23.4 | 20.2 | 17.0 |
| Tropical | Xishuangbanna1 | P3 | 16.7 | 18.3 | 21.1 | 23.6 | 25.3 | 25.8 | 25.2 | 25.3 | 24.7 | 23.1 | 19.9 | 16.7 |
| Tropical | Xishuangbanna2 | P1 | 16.4 | 17.8 | 20.4 | 23.0 | 24.7 | 25.2 | 24.8 | 24.8 | 24.2 | 22.7 | 19.6 | 16.4 |
| Tropical | Xishuangbanna2 | P2 | 16.4 | 17.8 | 20.4 | 23.0 | 24.7 | 25.2 | 24.8 | 24.8 | 24.2 | 22.7 | 19.6 | 16.4 |
| Tropical | Xishuangbanna2 | P3 | 16.4 | 17.8 | 20.4 | 23.0 | 24.7 | 25.2 | 24.8 | 24.8 | 24.2 | 22.7 | 19.6 | 16.4 |
| Tropical | Xishuangbanna3 | P1 | 16.3 | 18.0 | 20.9 | 23.1 | 24.6 | 25.2 | 24.6 | 24.7 | 24.1 | 22.5 | 19.4 | 16.2 |
| Tropical | Xishuangbanna3 | P2 | 16.3 | 18.0 | 20.9 | 23.1 | 24.6 | 25.2 | 24.6 | 24.7 | 24.1 | 22.5 | 19.4 | 16.2 |
| Tropical | Xishuangbanna3 | P3 | 16.3 | 18.0 | 20.9 | 23.1 | 24.6 | 25.2 | 24.6 | 24.7 | 24.1 | 22.5 | 19.4 | 16.2 |
| Tropical | Xishuangbanna4 | P1 | 16.1 | 17.8 | 20.6 | 22.8 | 24.2 | 24.8 | 24.2 | 24.3 | 23.8 | 22.2 | 19.1 | 16.0 |
| Tropical | Xishuangbanna4 | P2 | 16.1 | 17.8 | 20.6 | 22.8 | 24.2 | 24.8 | 24.2 | 24.3 | 23.8 | 22.2 | 19.1 | 16.0 |
| Tropical | Xishuangbanna4 | P3 | 16.1 | 17.8 | 20.6 | 22.8 | 24.2 | 24.8 | 24.2 | 24.3 | 23.8 | 22.2 | 19.1 | 16.0 |
| Tropical | Xishuangbanna5 | P1 | 17.1 | 18.7 | 21.5 | 24.1 | 25.8 | 26.4 | 25.7 | 25.8 | 25.2 | 23.5 | 20.3 | 17.1 |
| Tropical | Xishuangbanna5 | P2 | 17.1 | 18.7 | 21.5 | 24.1 | 25.8 | 26.4 | 25.7 | 25.8 | 25.2 | 23.5 | 20.3 | 17.1 |
| Tropical | Xishuangbanna5 | P3 | 17.0 | 18.6 | 21.4 | 23.9 | 25.6 | 26.2 | 25.6 | 25.6 | 25.1 | 23.4 | 20.2 | 17.0 |

**Table S7.** The mean monthly precipitation (mm) for the 39 sampling plots in temperate, warm-temperate, and tropical climatic zones.

| Climatic zone | Forest site | Plot | Jan | Feb | Mar | Apr | May | Jun | Jul | Aug | Sep | Oct | Nov | Dec |
| --- | --- | --- | --- | --- | --- | --- | --- | --- | --- | --- | --- | --- | --- | --- |
| Temperate | Liangshui1 | P1 | 3 | 4 | 9 | 24 | 57 | 100 | 163 | 156 | 73 | 29 | 11 | 6 |
| Temperate | Liangshui1 | P2 | 3 | 4 | 9 | 24 | 57 | 100 | 163 | 156 | 73 | 29 | 11 | 6 |
| Temperate | Liangshui1 | P3 | 3 | 4 | 9 | 24 | 57 | 100 | 163 | 156 | 73 | 29 | 11 | 6 |
| Temperate | Liangshui2 | P1 | 3 | 4 | 8 | 23 | 55 | 98 | 162 | 153 | 72 | 29 | 11 | 6 |
| Temperate | Liangshui2 | P2 | 3 | 4 | 8 | 23 | 55 | 98 | 162 | 153 | 72 | 29 | 11 | 6 |
| Temperate | Liangshui2 | P3 | 3 | 4 | 8 | 24 | 55 | 99 | 162 | 154 | 73 | 29 | 11 | 6 |
| Temperate | Changbaishan1 | P1 | 4 | 7 | 14 | 41 | 64 | 111 | 163 | 154 | 69 | 28 | 17 | 8 |
| Temperate | Changbaishan1 | P2 | 4 | 7 | 14 | 41 | 64 | 111 | 163 | 154 | 69 | 28 | 18 | 8 |
| Temperate | Changbaishan1 | P3 | 4 | 7 | 14 | 41 | 64 | 111 | 163 | 154 | 69 | 28 | 18 | 8 |
| Temperate | Changbaishan2 | P1 | 4 | 7 | 14 | 41 | 68 | 117 | 169 | 159 | 71 | 29 | 18 | 8 |
| Temperate | Changbaishan2 | P2 | 4 | 7 | 14 | 41 | 68 | 117 | 170 | 160 | 70 | 29 | 18 | 8 |
| Temperate | Changbaishan2 | P3 | 4 | 7 | 14 | 41 | 68 | 117 | 170 | 160 | 70 | 29 | 18 | 8 |
| Warm-temperate | Donglingshan | P1 | 3 | 5 | 11 | 21 | 47 | 78 | 145 | 113 | 50 | 20 | 9 | 2 |
| Warm-temperate | Donglingshan | P2 | 3 | 5 | 11 | 21 | 47 | 78 | 145 | 113 | 50 | 20 | 9 | 2 |
| Warm-temperate | Donglingshan | P3 | 3 | 5 | 11 | 21 | 47 | 78 | 145 | 113 | 50 | 20 | 9 | 2 |
| Warm-temperate | Jigongshan | P1 | 26 | 44 | 69 | 100 | 128 | 141 | 202 | 145 | 107 | 75 | 52 | 19 |
| Warm-temperate | Jigongshan | P2 | 26 | 44 | 70 | 102 | 130 | 143 | 204 | 146 | 108 | 76 | 52 | 20 |
| Warm-temperate | Jigongshan | P3 | 26 | 44 | 69 | 100 | 128 | 141 | 202 | 145 | 107 | 75 | 52 | 19 |
| Warm-temperate | Badagongshan | P1 | 32 | 42 | 81 | 153 | 205 | 231 | 216 | 182 | 154 | 121 | 68 | 33 |
| Warm-temperate | Badagongshan | P2 | 32 | 42 | 81 | 153 | 205 | 231 | 216 | 182 | 154 | 121 | 68 | 33 |
| Warm-temperate | Badagongshan | P3 | 32 | 42 | 81 | 153 | 205 | 231 | 216 | 182 | 154 | 121 | 68 | 33 |
| Warm-temperate | Tiantongshan | P1 | 53 | 74 | 108 | 122 | 144 | 184 | 119 | 160 | 177 | 80 | 65 | 45 |
| Warm-temperate | Tiantongshan | P2 | 53 | 74 | 108 | 122 | 144 | 184 | 119 | 160 | 177 | 80 | 65 | 45 |
| Warm-temperate | Tiantongshan | P3 | 53 | 74 | 108 | 122 | 144 | 184 | 119 | 160 | 177 | 80 | 65 | 45 |
| Tropical | Xishuangbanna1 | P1 | 17 | 26 | 30 | 63 | 169 | 189 | 298 | 277 | 274 | 153 | 110 | 42 |
| Tropical | Xishuangbanna1 | P2 | 17 | 26 | 30 | 63 | 169 | 189 | 298 | 277 | 274 | 153 | 110 | 42 |
| Tropical | Xishuangbanna1 | P3 | 17 | 25 | 30 | 62 | 166 | 183 | 292 | 270 | 279 | 157 | 112 | 42 |
| Tropical | Xishuangbanna2 | P1 | 16 | 29 | 28 | 64 | 168 | 188 | 297 | 281 | 315 | 163 | 112 | 37 |
| Tropical | Xishuangbanna2 | P2 | 16 | 29 | 28 | 64 | 168 | 188 | 297 | 281 | 315 | 163 | 112 | 37 |
| Tropical | Xishuangbanna2 | P3 | 16 | 29 | 28 | 64 | 168 | 188 | 297 | 281 | 315 | 163 | 112 | 37 |
| Tropical | Xishuangbanna3 | P1 | 18 | 25 | 29 | 60 | 167 | 184 | 293 | 273 | 269 | 157 | 106 | 40 |
| Tropical | Xishuangbanna3 | P2 | 18 | 25 | 29 | 60 | 167 | 184 | 293 | 273 | 269 | 157 | 106 | 40 |
| Tropical | Xishuangbanna3 | P3 | 18 | 25 | 29 | 60 | 167 | 184 | 293 | 273 | 269 | 157 | 106 | 40 |
| Tropical | Xishuangbanna4 | P1 | 18 | 25 | 28 | 59 | 166 | 184 | 292 | 274 | 266 | 157 | 104 | 38 |
| Tropical | Xishuangbanna4 | P2 | 18 | 25 | 28 | 59 | 166 | 184 | 292 | 274 | 266 | 157 | 104 | 38 |
| Tropical | Xishuangbanna4 | P3 | 18 | 25 | 28 | 59 | 166 | 184 | 292 | 274 | 266 | 157 | 104 | 38 |
| Tropical | Xishuangbanna5 | P1 | 18 | 26 | 31 | 64 | 170 | 191 | 300 | 281 | 273 | 153 | 108 | 43 |
| Tropical | Xishuangbanna5 | P2 | 18 | 26 | 31 | 64 | 170 | 191 | 300 | 281 | 273 | 153 | 108 | 43 |
| Tropical | Xishuangbanna5 | P3 | 17 | 26 | 30 | 63 | 169 | 189 | 298 | 278 | 275 | 154 | 110 | 43 |
